# binny: an automated binning algorithm to recover high-quality genomes from complex metagenomic datasets

**DOI:** 10.1101/2021.12.22.473795

**Authors:** Oskar Hickl, Pedro Queirós, Paul Wilmes, Patrick May, Anna Heintz-Buschart

## Abstract

The reconstruction of genomes is a critical step in genome-resolved metagenomics and for multi-omic data integration from microbial communities. Here, we present *binny*, a binning tool that produces complete and pure metagenome-assembled genomes (MAG) from both contiguous and highly fragmented genomes. Based on established metrics, *binny* outperforms or is highly competitive with commonly-used and state-of-the-art binning methods and finds unique genomes that could not be detected by other methods. *binny* uses k-mer-composition and coverage by metagenomic reads for iterative, non-linear dimension reduction of genomic signatures, as well as subsequent automated contig clustering with cluster assessment using lineage-specific marker gene sets. When compared to seven widely used binning algorithms, *binny* provides substantial amounts of uniquely identified MAGs and almost always recovers the most near-complete (>95% pure, >90% complete) and high-quality (>90% pure, >70% complete) genomes from simulated data sets from the Critical Assessment of Metagenome Interpretation (CAMI) initiative, as well as substantially more high-quality draft genomes, as defined by the Minimum Information about a Metagenome-Assembled Genome (MIMAG) standard, from a real-world benchmark comprised of metagenomes from various environments than any other tested method.

## 1 Introduction

High-throughput shotgun sequencing has become the standard to investigate metagenomes [1,2]. Metagenome-assembled genomes (MAGs) allow the linking of the genetic information at species or strain level: In the absence of cultured isolates, MAGs form an important point of reference. Thereby, study-specific MAGs have led to the discovery of previously uncharacterised microbial taxa [3] and deepened insights into microbial physiology and ecology [4,5]. In addition, large system-wide collections, which have been assembled recently, e.g. for the human microbiome [6] and several environmental systems [7], equip researchers with a common resource for short-read annotation. These collections also represent an overview of the pangenomic potential of microbial taxa of interest [8,9]. In addition to facilitating the interpretation of metagenomic data, genome-resolution also provides an anchor for the integration of functional omics [10, 11].

However, obtaining complete and un-contaminated MAGs is still challenging [12]. Most approaches start from assembled contigs, which are then binned by clustering, e.g. expectation-maximization clustering [13, 14] or graph-based clustering [15], of k-mer frequency or abundance profiles or both. Therefore, issues with metagenomic assemblies, such as fragmentation of the assembly because of insufficient sequencing depth, repeat elements within genomes, and unresolved ambiguities between closely related genomes, are perpetuated to MAGs. In addition, the features based on which contigs are binned are not generally homogeneous over genomes: for example copy number, and thereby metagenomic coverage, may vary over the replicating genome; certain conserved genomic regions, and also newly acquired genetic material, can deviate in their k-mer frequency from the rest of the genome [12].

In the face of these challenges, the algorithms used to bin assembled metagenomic contigs into congruent groups which form the basis for MAGs, can approximately be evaluated according to a set of criteria [16]. Most importantly, MAGs should be as complete as possible and contain as little contamination as possible. In metagenomic datasets with defined compositions, such as those provided by the Critical Assessment of Metagenome Interpretation (CAMI) initiative [17–19], the evaluation can be achieved by comparison with the reference genomes. For yet un-sequenced genomes, completeness and contamination can be assessed based on the presence and redundancy of genes that are expected to be present as single copies in many [20] or all [21] bacteria or archaea [22], or in specific lineages [23]. Contiguity and GC-skew provide further measures for highly complete genomes [12]. For reporting and storing MAGs in public repositories, the Minimum Information about a Metagenome-Assembled Genome (MIMAG) standard has been proposed [24]. In addition to completeness and contamination based on protein-coding genes, this standard also takes into account the presence of tRNA and rRNA genes. The latter present particular challenges for assembly and binning methods alike [12]. Nevertheless, the recruitment of rRNA genes to MAGs would improve the association with existing MAG collections [6,25] and rRNA-gene-based databases [26], which are widely used for microbial ecology surveys. In addition to binning tools, refiners have been developed that complement results from multiple binning methods [27,28]. These refiners generally improve the overall yield and quality of MAGs [29]. Finally, manual refinement of MAGs with the support of multiple tools is still recommended [12,30–33].

Here, we present *binny*, an automated binning method that was developed based on a semi-supervised binning strategy [10,34]. *binny* is implemented as a reproducible Python-based workflow using Snakemake [35]. *binny* is based on iterative clustering of dimension-reduced k-mer and abundance profiles of metagenomic contigs. It evaluates clusters based on the presence of lineage-specific single copy marker genes [23]. We benchmarked *binny* against six CAMI [17,18] data sets and compared the results with the most popular binning methods MetaBAT2 [15], MaxBin2 [14], CONCOCT [13], and the recently developed VAMB [36], SemiBin [37], and MetaDecoder [38]. We evaluated the contribution of *binny* to automatic MAG refinement using MetaWRAP [27] and DAS Tool [28]. Finally, we evaluated the MAGs returned by all approaches from real-world metagenomic datasets from a wide range of ecosystems. We report that *binny* outperforms or is highly competitive with existing methods in terms of completeness and purity and improves combined refinement results. *binny* also returned most MIMAG-standard high-quality draft genomes from both highly fragmented and more contiguous metagenomes over a range of microbial ecosystems.

## 2 Material and Methods

### *binny* workflow

*binny* is implemented as a Snakemake [35] workflow (Figure 1). At the centre of the workflow is the binning algorithm written in Python, which uses iterative, nonlinear dimension reduction of metagenomic read coverage depth and signatures of multiple k-mer sizes with subsequent automated contig clustering and cluster assessment by lineage-specific marker gene sets. Preparatory processing steps include the calculation of the average depth of coverage, gene calling using Prokka [39], masking of rRNA gene and CRISPR regions on input contigs, and identifying CheckM [23] marker genes using Mantis [40].

**Figure 1.**
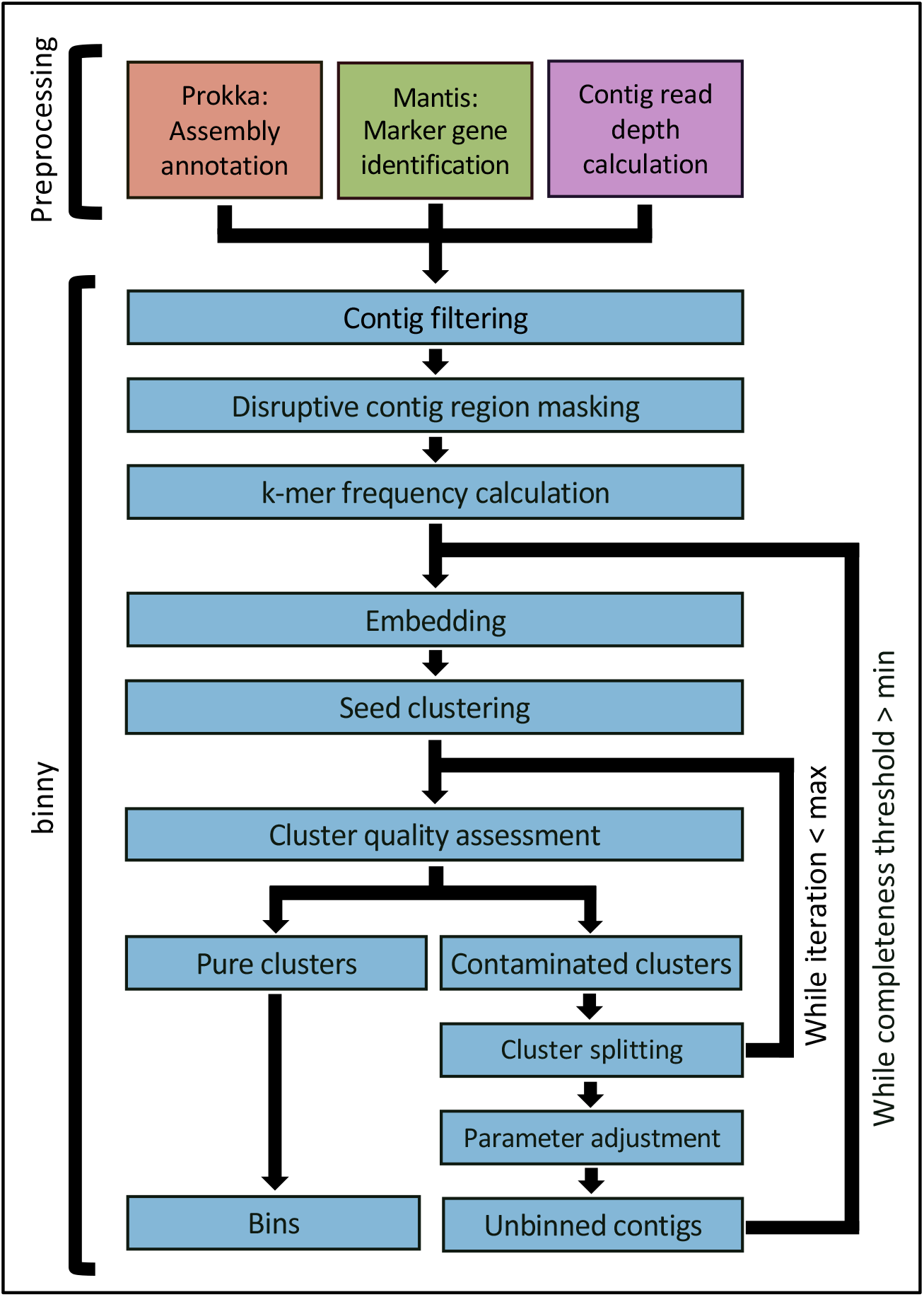
*binny* workflow. Overview of the Snakemake pipeline and of *binny*’s binning method. Preprocessing includes assembly annotation using Prokka, CheckM marker gene detection using Mantis, and (optional) average contig read coverage calculation. *binny* filters out contigs shorter than the specified value, masks potentially disruptive contig regions before calculating k-mer frequencies for the chosen k-mer size(s). In its main routine, *binny* iteratively embeds the contig data into two-dimensional space, forms clusters, assesses them with marker genes, and iteratively extracts clusters of sufficient quality as MAGs.

#### 2.0.1 Overview

*binny* operates in an iterative manner after processing of the annotated marker gene sets. Each iteration consists of non-linear dimension reduction on the selected features (depths of coverage and k-mer frequencies) of the so far unbinned contigs and clustering based on the resulting two-dimensional coordinates. Clusters are selected if the contained marker gene sets indicate purity and completeness above defined thresholds. A new iteration is started on left-over un-binned contigs with dynamically adjusted parameters. Finally, clusters above the thresholds are output as MAGs.

#### 2.0.2 Marker gene set processing

*binny* generates a directed graph database of the CheckM [23] taxon-specific marker sets annotated per contig in NetworkX [41]. This allows for fast access to the hierarchical (lineage-based) information. Some marker sets are omitted, as they are very small and/or led to imprecise assessments in testing (Supplementary Table 1).

#### 2.0.3 Filtering of short sequences

By default, *binny* filters out all sequences shorter than 500 bp. For its main routine, further filtering is done based on an Nx value (default 90). For Nx filtering, the contigs are sorted by length in descending order and the first contigs that together make up x% of the assembly are retained. This size selection can be modified by setting minimum size values or ranges for contigs that do not contain marker genes (default 2250 bp) and those that contain them (default 2250 bp). This aims to maintain the maximum amount of information from an assembly because only contigs that have a low information content are omitted.

#### 2.0.4 Masking of disruptive sequence regions

Specific regions on a sequence could skew the k-mer frequency and, thus, adversely affect the binning process. For example, CRISPR regions contain foreign genetic elements, which have k-mer frequencies that can deviate substantially from the rest of the genome, whereas rRNA genes have highly conserved sequences whose *k*-mer profiles do not resemble the rest of a given genome. To avoid an impact on the *k*-mer frequency calculation and still keep sequences intact, *binny* by default masks sequence elements/regions such as rRNA genes and CRISPR regions, using Prokka-provided annotations from barrnap [39] and minced [42], respectively. The masked regions are ignored during the *k*-mer frequency calculation.

#### 2.0.5 Single contig genome recovery

Genomes represented by single contigs might not be distinguishable from noise during clustering or be clustered together with highly similar contigs of other, fragmented genomes. Therefore, contigs with at least 40 different markers are extracted first and, if they are at least 90% pure and 92.5% complete, they are kept as single-contig MAGs and by default do not enter the iterative binning procedure.

#### 2.0.6 Binning features

*binny* uses two contig features for dimensionality reduction and clustering: the k-mer frequencies of multiple sizes (default k = 2, 3, and 4) and the average read coverage (raw read counts of one or more samples), both centered log-ratio transformed. Coverage information can be included in form of bam files or a file with tab-separated average contig coverage values per sample.

#### 2.0.7 Dimensionality reduction

To reduce the dimensionality of all features to two, the Fast Fourier Transform-accelerated Interpolation-based t-distributed Stochastic Neighbor Embedding implementation of openTSNE [43] is used. To decrease the computation time of the dimensionality reduction, Principal Component Analysis is used beforehand to lower the dimensionality of the initial feature matrix to either as many dimensions needed to explain 75% of the variation or to a maximum of 75 dimensions. To improve the embedding quality, especially with large datasets, multiple strategies are used: i) a multi-scale kernel with perplexity ranges from 10 to 20 and 100 to 130 starting with 10 and 100, where each iteration the former is increased by 2 and the latter by 5, are used instead of a Gaussian model to balance out local and global structure, as described by Kobak and Berens [44]. ii) An early exaggeration of *EX* for the number of unbinned contigs *NUC*:

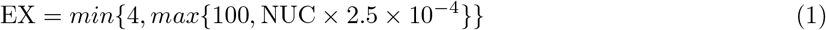

with a learning rate *LR_EX* for the early exaggeration phase:

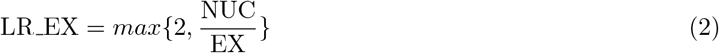

and a learning rate *LR*:

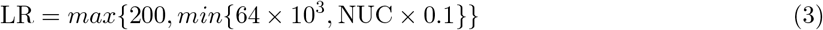

for the main phase are used. These values were chosen to achieve adequate embeddings of data sets of varying sizes [45,46]. Additionally, the number of iterations to run early and main phase optimizations are based on the difference in Kullback-Leibler divergence (*KLD*) *KLD_DIFF*. The KLD is measured every 250 optimization iterations. The optimization ends [46], if:

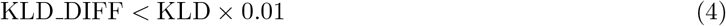

iii) To avoid the impreciseness of Euclidean distance measures in high dimensional space, Manhattan distance was chosen instead [47]. Default values were kept for all other openTSNE parameters.

#### 2.0.8 Iterative clustering

*binny* uses hierarchical density-based spatial clustering of applications with noise (HDBSCAN) [48] on the generated two dimensional embedding, in iterations. *binny* will run clustering of the created embedding n times (default 3), each time extracting MAGs meeting the quality thresholds and continuing with the embedding containing only the leftover contigs. n is the number of values for HDBSCAN’s `min_samples` parameter (default 1,5,10, hence n=3).

Other default clustering parameters are: the minimum cluster size is calculated with ln(n *contigs*), the cluster selection epsilon to merge micro-clusters is changed each *binny* iteration, cycling from 0.25 to 0.0 in 0.125 steps, and the distance metric used is Manhattan.

For each cluster, completeness and purity are assessed (see below). If a cluster passes the completeness threshold (by default starting with 92.5% and then decreasing to a minimum of 72.5%) and has a purity above 95%, if the completeness threshold is 90% or higher, otherwise it is set to 92.5%, it is kept as a MAG. Otherwise, *binny* will attempt to split that contig cluster iteratively using HDBSCAN a defined maximum amount of times (see above) but adding the raw depth(s) of coverage as additional dimension(s). Within each of these clustering rounds, the clusters below the quality threshold can be split again using HDBSCAN until no new clusters are identified and/or the maximum number of iterations is reached (default 1, no further splitting). To prevent the selection of low purity clusters, the purity threshold is increased continuously to a maximum of 95% at completeness 70% or lower (99%, if the chosen marker set is Bacteria or Archaea).

#### 2.0.9 Cluster assessment using marker gene sets

Clusters are assessed by calculating the purity and completeness based on the CheckM marker grouping approach, where marker genes known to be co-located in genomes of a lineage are collapsed into marker sets [23]. *binny* calculates MAG quality as in Parks et al. equation 1 and 2, respectively [23], except that instead of contamination purity is calculated: Let *P* be the purity for a set of collocated marker sets *MSS, MS* a marker set in *MSS, g* a single copy marker gene in MS, and *C* the counts of *g* in a MAG:

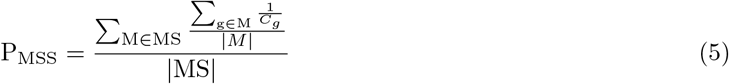

The taxonomic level and identity of the marker set are chosen dynamically: Assessment starts with completeness and purity of the domain-level marker sets and traverses the lineage down one taxonomic level at a time. At each level, completeness and purity for each taxon of the lineage are calculated. To combine the power of the domain level marker sets to give a general quality assessment with the specificity of lower level marker sets, the mean of purity and contamination for sub-domain level marker sets and their respective domain level set is used. If the marker set of the current taxon has an equal or higher completeness than the previously best-fitting marker set, it is set as the new reference. This choice is based on the assumption that the marker set with the highest completeness is least likely to be matching by chance and the larger the marker set size, the smaller the chance for miss-annotation. The lowest level to evaluate can be set by the user (default Class level).

#### 2.0.10 Iterative binning

*binny* starts embedding and clustering the size-selected, un-binned contigs. The minimum contig size limit is decreased by 500 bp if less than half of the iterative clustering steps returned MAGs, until a minimum size of 500 bp is reached. In the next binning iteration, the completeness threshold will be decreased by 10% and the initial contig size threshold reset to the initial maximum value after which the cycle starts again. This will continue until the minimum completeness threshold is reached. At this point, the purity threshold is decreased to 87.5% for clusters with completeness ≥ 90% and the number of splitting attempts for contaminated clusters is increased to 2. This is done to recover as much information as possible in the final binning iteration. *binny* has a separate routine for co-assemblies, i.e. runs with depth of coverage information from more than one sample: here, *binny* creates embeddings and clusters of the un-binned contigs ≥ 500 bp of and runs subsequent binning iterations, for as long as it finds new MAGs that satisfy the purity and completeness thresholds. The completeness threshold is decreased by 10% in every binning iteration, down to the minimum completeness threshold (default 70% completeness). As with the single sample mode, the purity threshold is decreased to 87.5% for clusters with completeness ≥ 90% and the number of splitting attempts for contaminated clusters is increased to 2. Once no more MAGs are found at the minimum completeness threshold, *binny* runs final rounds with minimum contig sizes starting at 2000 bp, decreasing by 500 each round, until 500 bp or the minimum size set by the user is reached.

#### 2.0.11 Contig depth of coverage calculation

If not provided explicitly, the average depth of coverage calculation can be performed directly from given BAM files within the Snakemake workflow using BEDTools [49] *genomeCoverageBed* and an in-house Perl script.

#### 2.0.12 Coding sequence, RNA gene, and CRISPR prediction by Prokka

A modified Prokka [39] executable is run with `--metagenome`, to retrieve open reading frame (ORF) predictions from Prodigal [50], rRNA and tRNA gene predictions from barrnap [39] and CRISPR region predictions from minced [42]. The modification improves speed by omitting the creation of a GenBank output and by the parallelization of the Prodigal ORF prediction step. Additionally, it allows the output of partial coding sequences without start- and/or stop-codons, which are frequently encountered in fragmented assemblies. No functional annotations of the called coding sequences are performed. The GFF output of Prokka is used in the subsequent steps.

#### 2.0.13 Marker gene set annotation

Taxon-specific marker gene sets are acquired from CheckM^1^ [23] upon installation of *binny*, hidden Markov profile models (HMM) of marker genes not found in `taxon_marker_sets.tsv` are removed, and `checkm.hmm` is split into PFAM [51] and TIGRFAM [52] parts. Mantis [40] is used to annotate coding sequences using the two HMM sets. Because both resources are of different scope and quality, consensus generation weights of 1.0 and 0.5 are used for PFAM and TIGRFAM models, respectively. Mantis’ heuristic search algorithm is used for hit processing, the e-value threshold is set to 1 × 10^-3^, and the `--no_taxonomy` flag is set.

#### 2.0.14 Parameter customization

To optimize for their use case, a user can choose to change the sizes and number of *k* -mers used, the Nx value and/or minimum contig length to filter the assembly, as well as the minimum completeness and purity thresholds for MAGs. The user may choose not to mask potentially disruptive regions and can control the clustering process by adjusting several HDBSCAN parameters. Additionally, it is possible to choose between internal calculation of the average contig read depth or supplying a depth value file.

#### 2.0.15 Requirements/dependencies

*binny* is implemented as a Snakemake pipeline and an installation script is provided that takes care of the installation of all necessary dependencies and required databases.

### Benchmarking

#### 2.0.16 Synthetic benchmark data

Binning performance was evaluated using data sets from the CAMI initiative [17, 19], each containing several hundreds of genomes at strain-level diversity. To benchmark against data of varying complexity, five short-read data sets with a total of 49 samples were chosen from the 2nd CAMI Toy Human Microbiome Project Dataset^2^. Additionally, to test against a very large data set, the five sample Toy Test Dataset High Complexity from the first CAMI challenge^3^ was used.

To test the performance on co-assembled data, the pooled assemblies of each of the six CAMI datasets and the respective number of sample read files for each data set, provided by the CAMI challenges, were used. Contig read depth per sample was calculated using *binny* and provided to all binning methods unless stated otherwise. Read files were de-interleaved^4^ and mapped against the contigs using `bwa-mem` [53].

#### 2.0.17 Real world benchmark data

To assess the binning performance in different real-world scenarios with a variety of metagenome sizes, complexities and qualities, 105 metagenomes used in the MetaBAT2 publication [15] for benchmarking were chosen based on the availability of preprocessed read data at the Joint Genome Institute (JGI). The newest available assembly for the metagenomes and the respective preprocessed reads were retrieved from JGI^5^. The read data were processed in the same way as the CAMI data. For a full list with all sample information see Supplementary Table 2.

#### 2.0.18 Binning and refinement methods

The performance of *binny* was compared with six other state-of-the-art binning methods, and to two binning refinement tools. *binny* and the other methods were all run using the default settings, unless specified otherwise:

MaxBin2 (2.2.7) [14] was run by providing the contig read depth files using the `-abund` option, and with the `-verbose` option.

MetaBAT2 (2.2.15) [15] was provided the contig read depth files using the `-a` option, and the options `-cvExt, --saveCls`, as well as `-v`.

CONCOCT (1.1.0) [13] was run following the ‘Basic Usage’ section in the documentation^6^.

VAMB (3.0.2) [36] was run with the default parameters and using the Snakemake pipeline as described in the documentation^7^. Because VAMB is designed to achieve optimal performance through the combination of the data of multiple samples, the samples from each of the six CAMI data sets were concatenated and run together, as described by the authors (README sections `Recommended workflow` and `Snakemake workflow`). For the real-world metagenomes, samples sharing a JGI GOLD Study ID were run together. As VAMB could not be successfully run on some of the real-world samples using default values, or when trying with lower values of `-m` and `--minfasta`, the number of MAGs recovered was counted as zero for these samples. For a list of these samples see Supplementary Table 3.

SemiBin (1.0.2) [37] was run using the `single_easy_bin` mode with `--random-seed 0` and default parameters otherwise. For the single sample binning the global model was used, except for the CAMI 2 Gastrointestinal tract samples, for which `--environment human_gut` was used and the CAMI 2 Oral samples, for which `--environment human_oral` was used. For the real-world benchmark the respective models matching wastewater, ocean, and soil samples were employed.

MetaDecoder (1.0.9) [38] was run using the default parameters, following the developers instructions, calling consecutively `coverage, seed` and `cluster`. To use `coverage`, the assemblies’ respective bam files were converted to sam format using samtools.

DAS Tool (1.1.2) [28] was run using Diamond [54] as a search engine on the unfiltered binning method outputs.

MetaWRAP (1.2.2) [27] was set to output only contigs with less than 10% contamination and at least 70% completeness and was also provided the unfiltered binning method outputs. Both refinement tools, DAS Tool and MetaWRAP, were run: i) per sample using the data of binny, MetaDecoder, and SemiBin and ii) the two binning methods except *binny*, to asses how many MAGs *binny* contributes in an ensemble approach.

#### 2.0.19 MAG quality standards

To match real-world workflows, all binning outputs were assessed using CheckM (1.0.12) [23] and filtered to contain only MAGs with a purity > 90% and a completeness > 70%. The latter threshold was set in accordance with the CheckM publication, which suggests that CheckM results are reliable at completeness equal or larger than 70%. MAGs above these thresholds are subsequently called ‘high quality’ (HQ) MAGs. MAGs with a purity > 95% and a completeness > 90% are called ‘near complete’ (NC) MAGs, as defined by Bowers et al. [24].

Additionally, the MIMAG definition of high-quality draft genomes was employed, requiring at least 18 unique tRNAs and three unique rRNAs to be present in the MAG in addition to a purity of > 95% and a completeness of > 90% [24].

Besides the recall in terms of bps of the assembly recovered, the read recruitment of MAGs was assessed. All reads mapping as primary mappings to contigs of a MAG were counted per sample and divided by the total read count (forward + reverse) using pysam^8^.

#### 2.0.20 Assessment of benchmark results

Results for the CAMI benchmark were processed using AMBER (2.0.3) [55], a genome reconstruction evaluation tool, with the following parameters, `-x ‘50,70,90’` and `-k ‘circular element’`.

To evaluate a MAG, AMBER selects the gold standard genome with the highest share of bps in that MAG as the reference. In contrast to CheckM, where purity and completeness refer to the amount of marker genes present or duplicated, within AMBER and using an available gold standard, purity and completeness refer to the amount of bp of the reference genome recovered for completeness, and the share of bp of a given MAG with a given reference genome, respectively. Additionally, to assess one or multiple data sets taken together, AMBER defines overall completeness as *‘Sum of base pairs coming from the most abundant genome in each predicted genome bin divided by the sum of base pairs in all predicted bins*.…’ and overall purity as ‘Sum *of base pairs coming from the most abundant genome in each predicted genome MAG divided by the sum of base pairs in all predicted bins*. …’.

Purity and completeness values are reported as the per data set average, unless specified otherwise. For the real-world benchmarks, the average proportion of bp recovered or the number of MAGs recovered is reported together with the standard error of the mean (SEM). Another metric used is the adjusted Rand index (ARI), which is a commonly used metric to measure how similar two datasets are. Trying to make the comparisons between different binning methods as realistic, fair and transparent as possible, we report all metrics derived from the CheckM-filtered binning results, unless specified otherwise.

To assess the intersections of MAGs formed by the different binning methods on multi-sample datasets, genomes were counted separately for each sample. To this end, the gold standard genome name was concatenated with the sample id to yield unique identifiers for each genome in each sample. All other figures were created using the Python libraries *matplotlib* [56] and *Seaborn* [57], as well as *UpSetPlot* [58], setting the minimum intersection size to be shown to ten, unless specified otherwise, for the UpSet plots. The remaining data analyses were performed and table outputs created using the Python *NumPy* and *pandas* libraries.

To evaluate if the binning methods could recover NC and HQ MAGs from organisms with closely related or highly similar genomes in the same sample, for each of the 54 samples of the six CAMI data sets all versus all Average Nucleotide Identity (ANI) calculations were performed using FastANI (1.33) [?]. Each genome was assigned the highest ANI to another genome in the same sample. The numbers of NC and HQ MAGs recovered per binning method with ANIs higher than 90.0-99.9% in 0.1 steps were counted.

## Results

### Performance on synthetic data sets

To assess *binny*’s performance, six datasets from the CAMI initiative were chosen: the high complexity toy dataset of the first CAMI iteration to investigate how *binny* performs on very large data sets and the five toy human microbiome data sets of the second CAMI iteration to evaluate the performance on a wide range of microbiome sizes and complexities. Generally, a binning tool performs best, if it recovers the most complete MAGs with the highest purity, which corresponds to the highest ARI.

Over all six data sets (54 samples), *binny* with default settings recovered 35.5% (SEM 2.8%) of the reference genome lengths in the samples as NC MAGs (n=1564) and 42.7% (SEM 3.0%) as HQ MAGs (n=2021), with median recall values of 26.3% and 36.3%, respectively (Figure 2, Supplementary Table 4). In total, 45.1% of the reference genomes where recovered at a purity of 98.4% with an ARI of 0.977 (Supplementary Figure 1, Supplementary Table 5).

**Figure 2.**
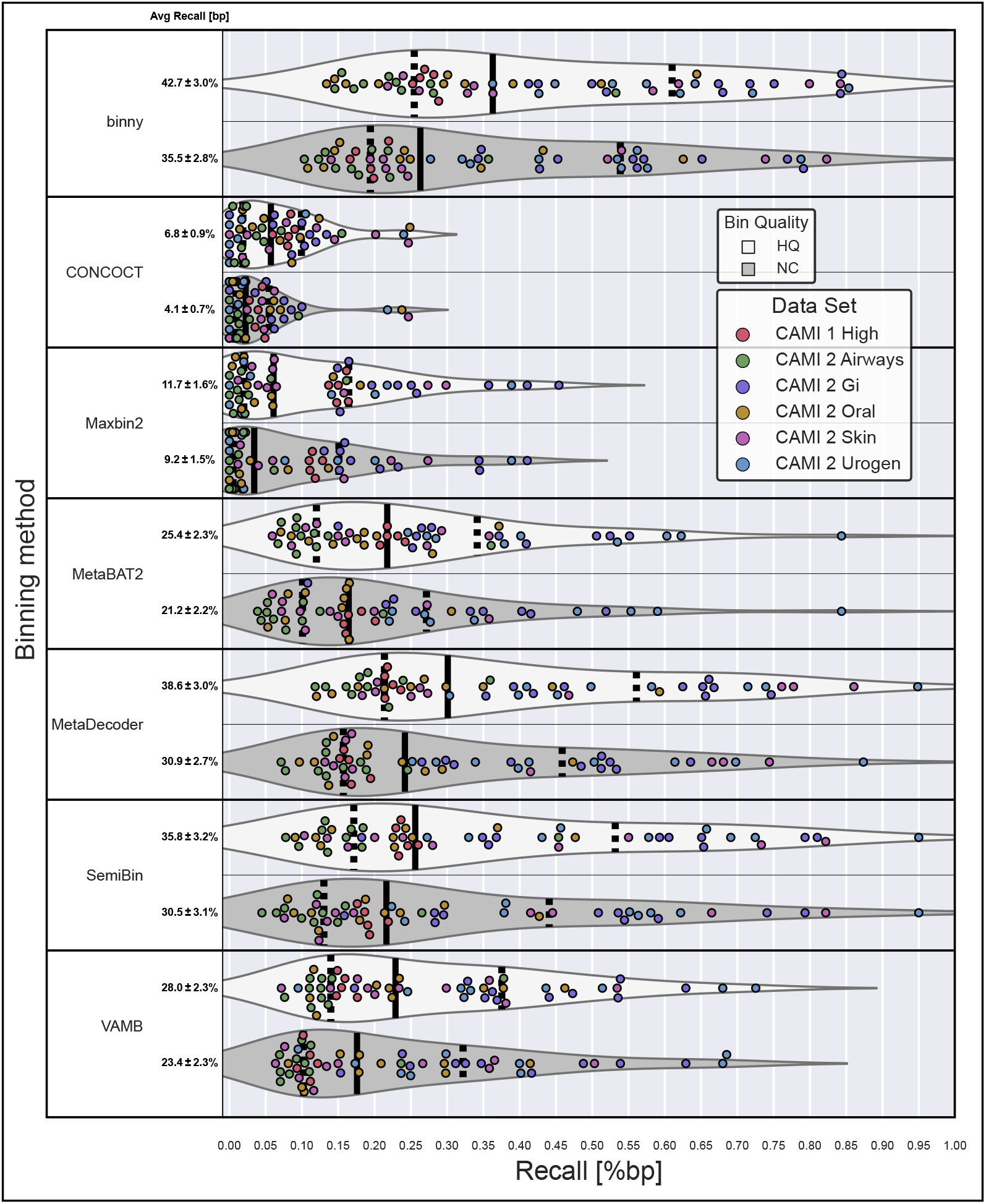
Performance of binning methods on CAMI data sets. Recall of bp assembled sequences as high-quality (HQ) and near-complete (NC) MAGs per binning method per sample, for the six CAMI data sets. The average recall is shown with the SEM.

For the high complexity data set, *binny* recovered 30.0% of the total reference genomes with a purity of 97.8% and an ARI of 0.970 (Supplementary Figure 2, Supplementary Table 6).

The lowest recall was observed for the CAMI 2 Airways data set with 25.9%, a purity of 98.1%, and an ARI of 0.973 (Supplementary Figure 3), whereas the highest recall of 66.3%, with a purity of 98.6% and an ARI of 0.978 was reached with the CAMI 2 GI data set (Supplementary Figure 4). For the other three data sets, *binny* achieved the following respective recall, purity and ARI numbers: 60.9%, 98.0% and 0.969 (CAMI 2 Urogenital); 48.0%, 98.9% and 0.983 (CAMI 2 Skin); and 33.2%, 98.6%, and 0.982 (CAMI 2 Oral) (Supplementary Figures 5-7, Supplementary Table 6; for detailed metrics for MAGs and samples see Supplementary Tables 7 and 8, respectively).

The average read recruitment from the CAMI data of the *binny* output was 72.4%. The highest recruitment was achieved for the GI data set sample 5 with 99.4%, whereas the lowest was observed for the skin dataset sample 19 (40.7%). Notably, a substantial proportion of the reads recruited were mapped to single contig MAGs for the CAMI 2 datasets (on average 60.7%), whereas for the CAMI 1 datasets, only about a fifth of the reads recruited by binned contigs, were mapped to single contig MAGs (Supplementary Tables 9 and 10).

#### 2.1 **Running** *binny* with multiple depth files

When assessing the performance on co-assembled datasets with depth information from multiple samples, *binny* had a recall of 54.3% over the CAMI datasets with a purity of 98.4%. In total 1055 NC MAGs were produced, 413 of which contained more than five contigs (Supplementary Figures 8-10). The highest recall was achieved for the CAMI 2 Gastrointestinal co-assembly with 75.9% and a purity of 99.0%, whereas the worst performance was observed for the CAMI 2 Airways dataset with a recall of 32.6% and purity of 97.4% (Supplementary Tables 11-13).

To test to which degree *binny* makes use of the information from the multiple read depth files per co-assembly, *binny* was additionally run with only one depth file per co-assembly. *binny* using all available depth files had a 20.4% higher recall at a slightly higher purity, leading to a recovery of 25.0% more NC MAGs (211) in total and 102.5% more NC MAGs (209) of contig sizes larger than 5 (Supplementary Figures 8-10, Supplementary Tables 11-13).

#### 2.2 Effect of masking potentially disruptive sequence regions

To test the effect of masking potentially disruptive sequences, we also ran *binny* on the 54 CAMI samples without the masking procedure. The unmasked run did not differ substantially from the one with the default settings regarding assembly recall and purity (Supplementary Table 14). In total, 29 fewer NC MAGs were recovered without the default masking (Supplementary Table 15). The amount of MAGs recovered matching the MIMAG standard was reduced by 5% from 1167 to 1112 (Supplementary Table 16).

#### 2.3 Effect of lineage-specific marker gene sets

To evaluate the utility of using lower taxonomic level marker gene sets, we compared the difference in NC and HQ MAGs recovered between the default setting of a maximum depth at class-level to only using kingdom-level markers with the unfiltered output from the 54 CAMI samples. With the class-level marker sets and 8.5% more NC and 21.0% more HQ MAGs with a size of more than five contigs could be recovered, demonstrating the effectiveness of the lower level marker gene information with *binny*. Overall, with class-level markers the recall was 5.7% higher, whereas the purity was 1.2% and the ARI 1.8% lower (Supplementary Tables 17 and 18).

#### 2.4 Run time

For all experiments, *binny* was run on compute nodes equipped with AMD Epyc ROME 7H12 CPUs, and for the run-time benchmark 32 cores and 56 GB of RAM were used. For the CAMI samples, the complete *binny* pipeline took on average 112 minutes to run, with a max of 413 minutes for sample five of the CAMI 1 high complexity data set. The Prokka annotations took on average 28%, the Mantis annotations on average 15% and *binny* on average 57% of the total run time (Supplementary Table 19).

### *binny* generally outperformed state-of-the-art binning methods on synthetic data

Over all six CAMI datasets *binny* recovered per sample the highest portion of the assembly (bps) as HQ (42.7%) or NC (35.5%) MAGs, followed by MetaDecoder (38.6%, 30.9%) and SemiBin (35.8%, 30.5%). Additionally, *binny* showed the highest median MAG counts with 23.8%, 36.8% more NC and 14.8%, 29.2% more HQ MAGs than MetaDecoder and SemiBin, respectively (Figure 2, Supplementary Table 4).

*binny* was the only binning method that resulted in high purity (97.3%) and high ARI (0.962) output over all datasets without additional CheckM filtering. Using CheckM filtering, *binny*’s purity and ARI were increased by 1.1% and 0.015, respectively, whereas the assembly recall was decreased by 3.0% (Supplementary Figure 1 b,c, Supplementary Table 5). The binning method with the second highest NC MAG recall, MetaDecoder, had a purity of 84.6% natively and an ARI of 0.813. After CheckM filtering, the purity and ARI of VAMB was the highest among binning methods (99.5% purity and an ARI of 0.994, respectively), but at the same time the recall was reduced from 56.7% to 28.5% (Supplementary Figure 1 b,c, Supplementary Table 5). For detailed metrics on the MAGs and samples see Supplementary Tables 7 and 8, respectively.

*binny* also outperformed the other binning methods on each of the individual data sets, except for the CAMI 1 High complexity dataset, where SemiBin produced 2.4% more NC MAGs (Figure 2, Supplementary Figure 2-7, Supplementary Table 7).

Many of the CAMI samples contain larger amounts of single-contig or almost contiguous genomes than are commonly observed in real-world samples. To evaluate *binny*’s performance without those, we considered the subset of genomes that consisted of more than five contigs. Here, *binny* also produced substantially more NC (13.1%) and HQ (25.3%) MAGs than the second best performing method, SemiBin (Supplementary Figure 11). *binny* recovered the largest amount of NC MAGs for the CAMI 2 GI, AW and Skin datasets, tied with SemiBin for the UG data set and came second for the Oral data set after VAMB (5.6% less) and the CAMI 1 High complexity data set after SemiBin (0.4% less), respectively (Supplementary Figure 12, Supplementary Table 7). Looking at the assembly recall as NC and HQ MAGs, *binny* showed the best performance for all data sets (Supplementary Figure 13).

Additionally, *binny* recovered the most NC and HQ MAGs on co-assembly versions of the six data sets. It recovered 9.2% more NC and 13.9% more HQ MAGs than the second best method, MetaDecoder, and 7.6% more NC and 25.1% more HQ MAGs of genomes consisting of more than five contigs than the second best performer, SemiBin (Supplementary Figures 9, 10, 14, 15, Supplementary Table 11-13).

Lastly, we assessed the amount of MAGs meeting the MIMAG draft standard. *binny* recovered the most MAGs of that quality for each CAMI data set, recovering in total 20.3% more, with 1167, than the second best method, MetaDecoder, which produced 971 MIMAG drafts over all 54 samples from the six CAMI data sets(Table 1 and Supplementary Table 16).

**Table 1.** MAGs matching the MIMAG standard. Rows represent values per binning method for the six CAMI data sets and the number for the real-world benchmark data. Bold values show the highest count per data set, underlined values the second highest count.

|  | High | Airways | GI | Oral | Skin | Urogen | Total | IMG |
| --- | --- | --- | --- | --- | --- | --- | --- | --- |
| **binny** | **164** | **192** | **202** | **243** | **215** | **152** | **1168** | **629** |
| CONCOCT | 17 | 10 | 19 | 37 | 18 | 6 | 107 | 142 |
| MaxBin2 | 85 | 5 | 79 | 20 | 26 | 16 | 231 | 422 |
| MetaBAT2 | 144 | 81 | 100 | 134 | 85 | 111 | 655 | 417 |
| MetaDecoder | 140 | <u>147</u> | 166 | 193 | <u>181</u> | <u>144</u> | <u>971</u> | 533 |
| SemiBin | <u>148</u> | 113 | <u>168</u> | 184 | 156 | 143 | 912 | <u>553</u> |
| VAMB | 107 | 121 | 138 | <u>197</u> | 123 | 113 | 799 | 406 |

### *binny* recovered unique MAGs

To evaluate the performance of different binning tools, it is also of interest to see how much unique information is recovered by each individual binning method. *binny* yielded 42.5% more unique NC MAGs (114) than the next best, VAMB for the CAMI data sets. Additionally, the two largest sets of MAGs shared by two binning methods are both *binny* sharing MAGs with MetaDecoder (140) or SemiBin (57), respectively (Figure 3). For the HQ genomes, similar results were observed: *binny* recovered the second most unique MAGs after VAMB (5.8% less) and was present in all of the intersections with the largest numbers of genomes (Supplementary Figure 16, Supplementary Table 7). On the co-assemblies, *binny* recovered 31.3% more unique NC and 67.4% more unique HQ MAGs, than the method with the second most unique MAGs, MetaDecoder (Supplementary Figures 9 and 14).

**Figure 3.**
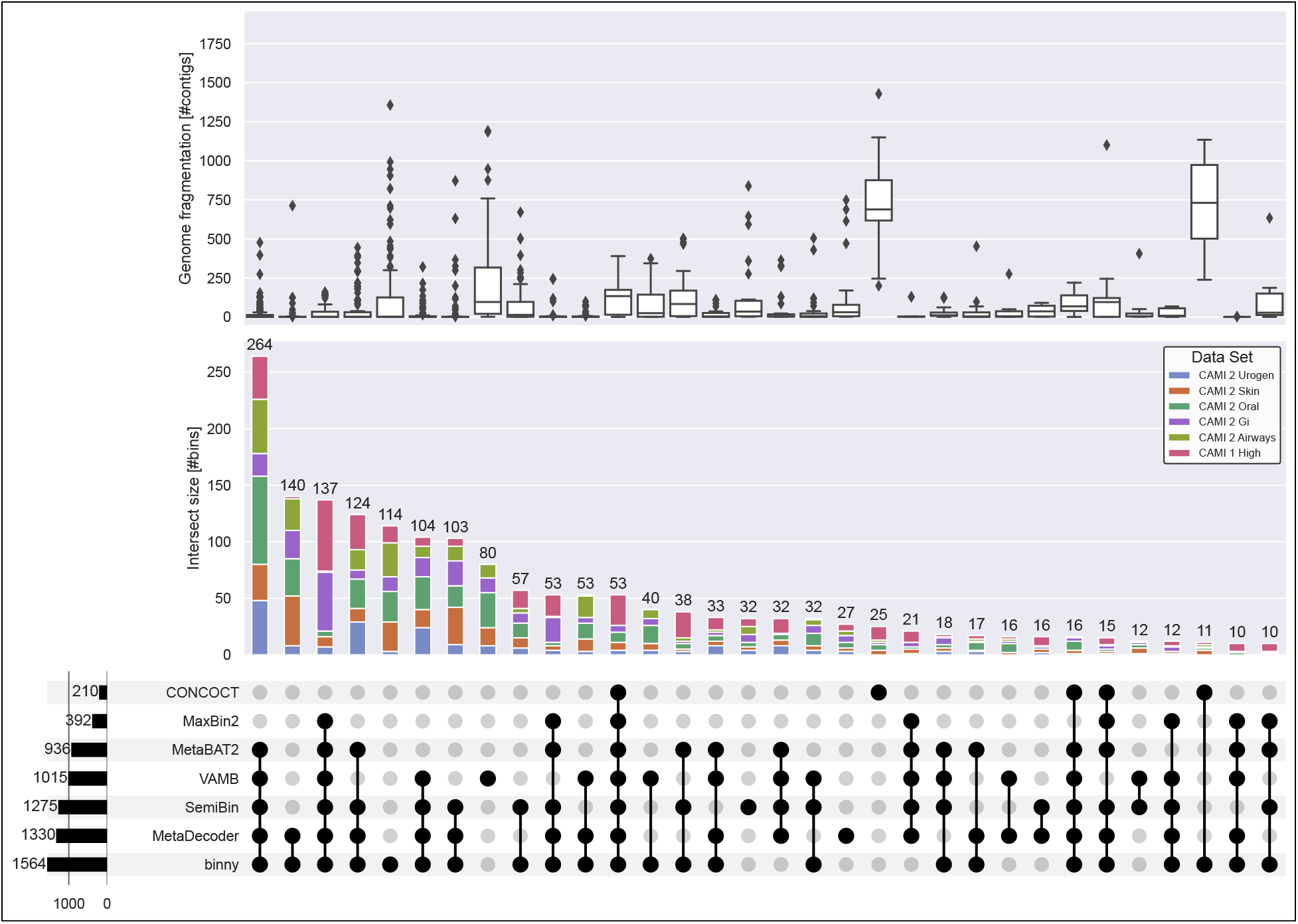
Intersections of recovered CAMI NC MAGs and reference genome fragmentation grade. Intersections of NC MAGs of seven CheckM-filtered binning methods for 54 samples from six CAMI data sets. Upper panel: Reference genome fragmentation in number of contigs. Middle panel: Intersection size in number of NC MAGs with proportions of MAGs stemming from the six CAMI data sets. Lower panel: Number of MAGs per binning method on the left, intersections > 9 in the centre.

### *binny* produced complete and pure MAGs from contiguous as well as highly fragmented genomes

Next, we assessed the ability of different binning methods to recover genomes of different fragmentation grades. *binny* recovered substantially more highly fragmented genomes (defined here as genomes with more than 500 contigs) than almost all methods (50 NC MAGs). Only CONCOCT recovered more highly fragmented genomes than *binny* (54), whereas both methods shared the recovery of a large portion of these fragmented genomes. VAMB produced the third most with 27 highly fragmented NC MAGs (Supplementary Figure 17a, Supplementary Table 7). When looking at the number of fragmented HQ MAGs recovered, *binny* substantially outperformed all other methods, recovering 26.6% more than the second best method, CONCOCT, with 282 MAGs (Supplementary Figure 17b, Supplementary Table 7). For the co-assemblies, *binny* recovered 133.3% more NC and 101.2% more HQ MAGs than the second best method SemiBin (Supplementary Figure 18, Supplementary Table 13).

#### *binny* recovers MAGs from genomes with highly similar relatives

When assessing a binning methods’ performance, it is also of interest to evaluate how well it is able to separate closely related organisms, as this would e.g., allow for the study of strain variation within a sample. Over all CAMI samples, *binny* recovered the largest amount of NC and HQ MAGs from genomes with highly similar relatives in the same sample over an ANI range from 90% to 99.9%. At an ANI of 90.0% binny recovered 730 NC and 840 HQ MAGs. The second and third highest performing methods were MetaDecoder (35.4% less NC, 20.2% less HQ MAGs) and SemiBin (52.1% less NC, 45.6% less HQ MAGs). When taking only into account genomes consisting of six or more contigs, *binny* still outperformed all other methods, followed by SemiBin (21.7% less NC, 19.8% less HQ MAGs) and VAMB (60.0% less NC, 25.5% less HQ MAGs). At an ANI cut-off of 95% *binny* recovered 36.8% and 22.6% more NC MAGs than the second highest performing method from genomes consisting of any number or at least six contigs, respectively. Finally, at an ANI of 99.0% *binny* was able to generate 41.8% and 61.1% more NC MAGs from genomes consisting of any number or at least six contigs, respectively, than the method placing second(Figure 4, Supplementary Figure 19).

**Figure 4.**
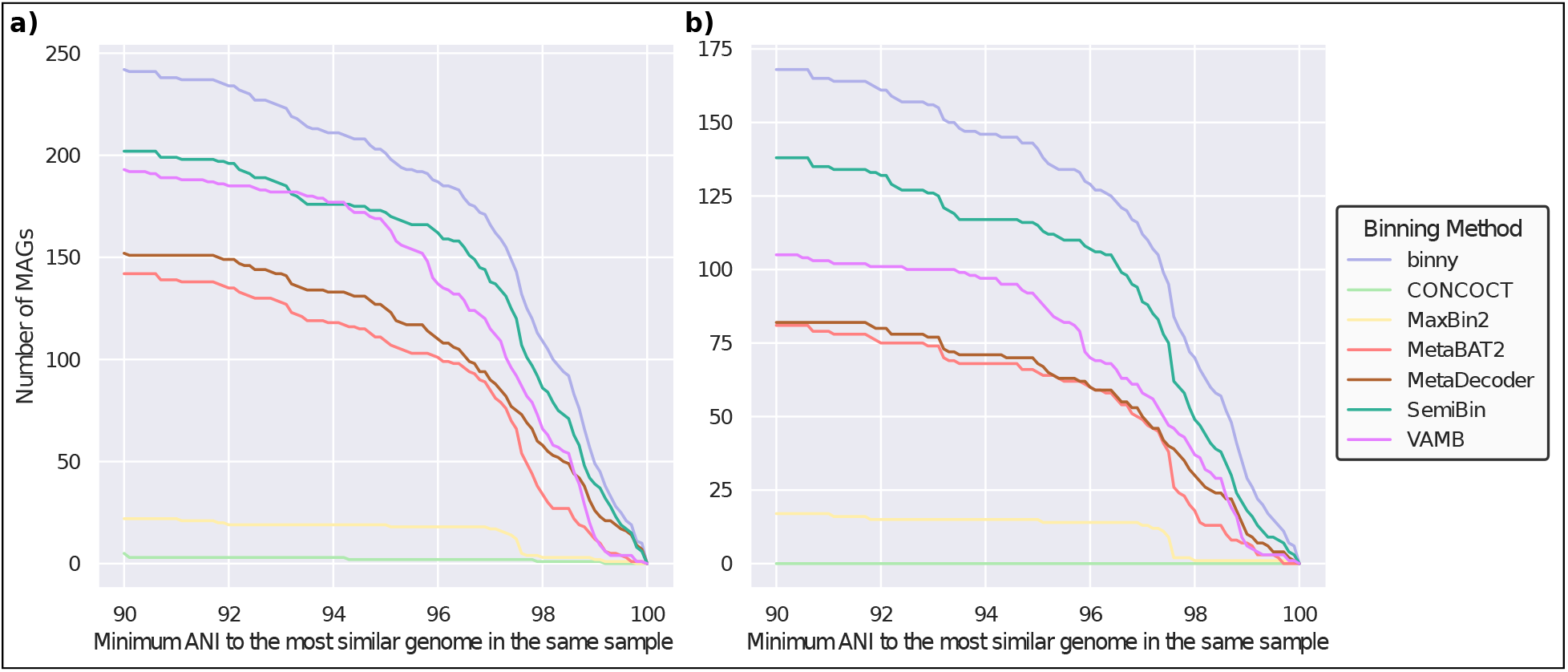
Performance of binning methods on recovering MAGs with close relatives. Number of a) HQ, and b) NC MAGs with a minimum ANI to the most similar genome in the same CAMI sample of at least 90.0% up to 99.9% in 0.1% steps for seven CheckM-filtered binning methods. Includes genomes consisting of at least six contigs.

### *binny* recovered the largest number of MIMAG drafts for real-world assemblies from different environments

When benchmarking binning tools with real-world data from a wide variety of environments, *binny* recovered on average the second largest amount of the assembly (bp) as NC (19.8%) bins, after MetaDecoder (20.2%), and the largest amount of HQ (28.8%) MAGs. MetaDecoder in total recovered the most NC MAGs (1647), followed by *binny* (1523) and SemiBin (1513). Notably, there was a substantial gap in performance to the next best method, MetaBAT2, with 1223 NC MAGs recovered (23.7% less than SemiBin). *binny* recovered the largest amount of HQ MAGs (3013), followed by MetaDecoder (2969) and SemiBin (2747). As in the CAMI benchmarks, CONCOCT showed the lowest recall for both NC and HQ MAGs, whereas MaxBin2 performed comparatively better with this data than in the CAMI benchmark (Figure 5 and Supplementary Tables 20, 21). When counting the recovered MAGS matching the MIMAG standard, *binny* produced 629 MAGs, 13.7% more than the second best performing method, SemiBin (553), with MetaDecoder ranking third with 533 (Table 1 and Supplementary Table 22).

**Figure 5.**
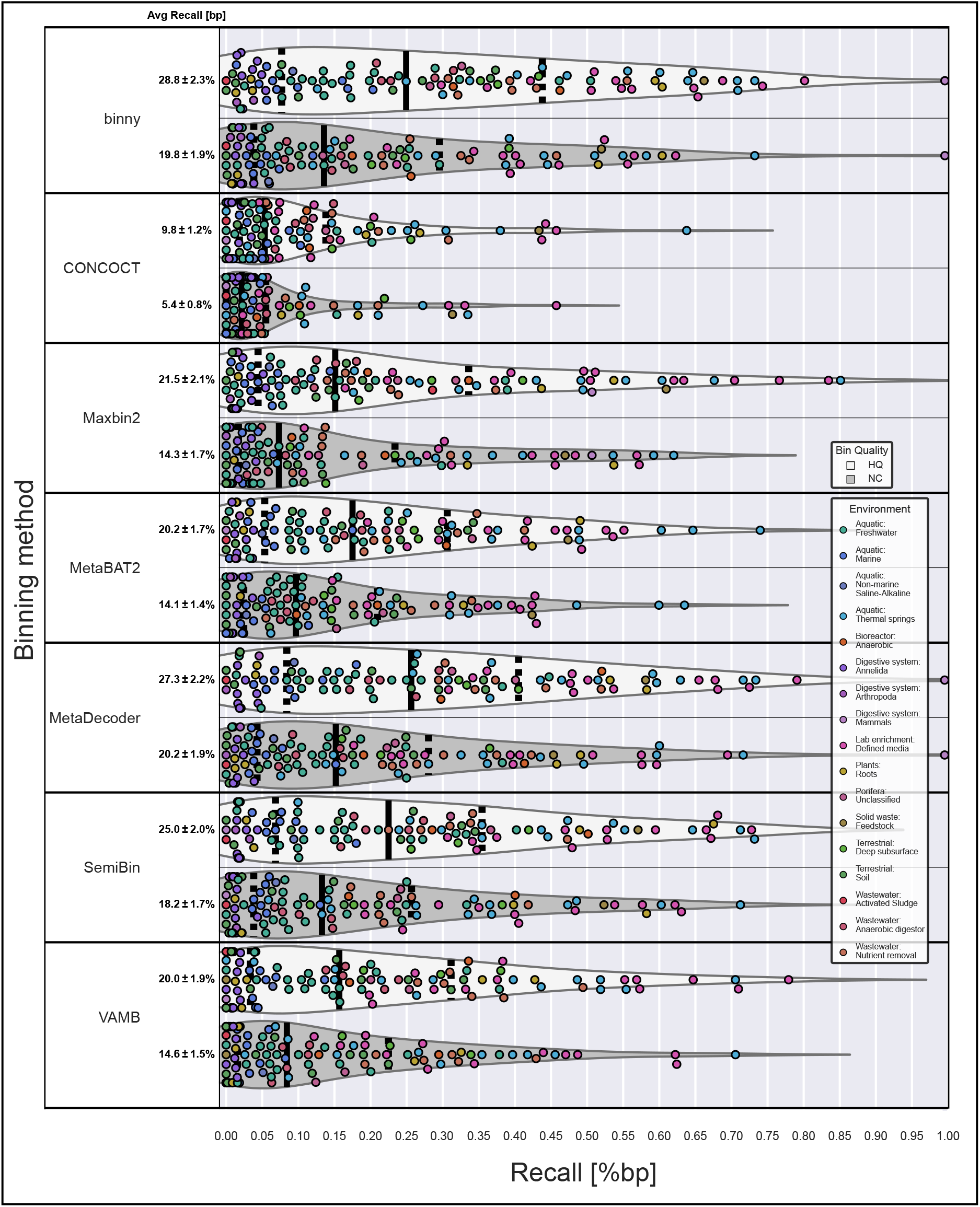
Performance of binning methods on real-world data sets from various environments. Assembly recovery as HQ and NC MAGs per binning method per sample from 105 real-world samples. The average recall (% bp) is shown with the SEM.

### *binny* improved ensemble binning/refinement approaches

To test if *binny* is able to improve refinements in combination with other binning methods, we ran the two most popular automatic refinement tools, DAS Tool and MetaWRAP, on the 54 samples of the six CAMI data sets, combining MetaDecoder and SemiBin either with or without *binny*.

When *binny* was excluded, a 1.9% and 2.9% lower recall was observed for DAS Tool (48.4%) and MetaWRAP (45.0%), respectively, whereas binny had marginally lower recall than DAS Tool with 48.1% (Supplementary Figure 20 b,c, Supplementary Table 23). *binny* on its own, unfiltered, recovered 7.0% more NC MAGs than DAS Tool and 2.4% less than MetaWrap without the *binny* MAGs. When including *binny*, MetaWRAP was able to recover 8.8% more NC MAGs (1705) than *binny* on its own, whereas DAS Tool produced 2.4% more NC MAGs (1605) (Supplementary Figure 20 a, Supplementary Figure 21, Supplementary Table 24). Only MetaWrap with *binny* input produced more HQ MAGs than *binny* alone with 2174 (6.0% more) (Supplementary Figure 22, Supplementary Table 24). Including only MAGs with more than five contigs, the DAS Tool and MetaWRAP without *binny* performed worse than *binny* alone (10.8% and 5.5% fewer NC MAGs, respectively). The runs including all three binning methods showed the highest performance overall, with MetaWRAP recovering the most MAGs (Figure 6). Evaluating the HQ MAG recovery the results were similar, but now only MetaWrap with all three binning methods outperformed *binny* (Supplementary Figure 23). While MetaWRAP produced almost no heavily contaminated MAGs, DAS Tool returned large numbers of MAGs with very low purity, despite showing over the entire CAMI benchmark data high purity (Supplementary Figure 20 d, Supplementary Table 23).

**Figure 6.**
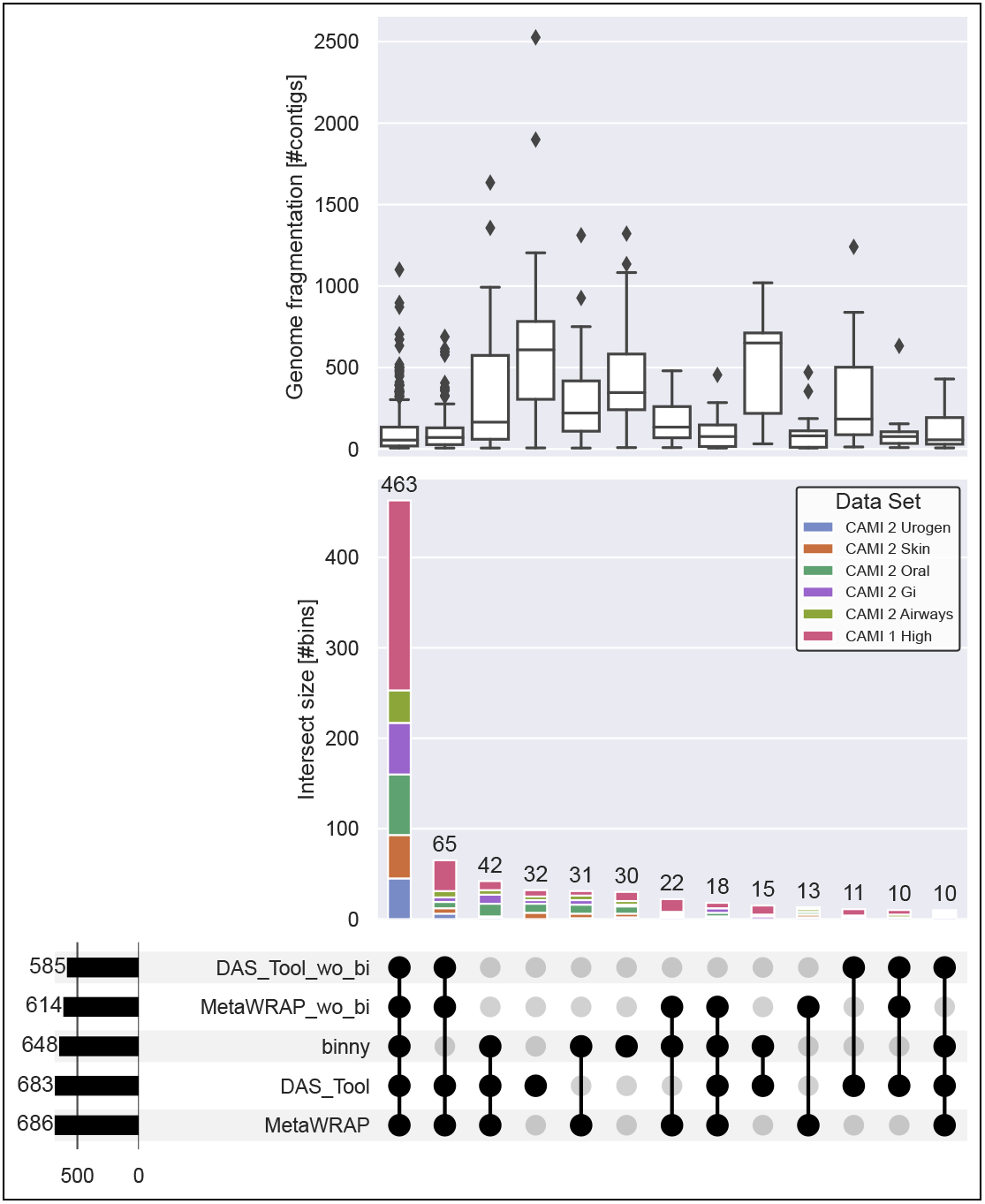
Intersections of recovered CAMI NC MAGs from bin refinement methods. Intersections of NC (near-complete) MAGs from genomes consisting of more than five contigs by *binny*, DAS Tool and MetaWrap for 54 samples from six CAMI data sets. Binning method output used by the refinement methods: *binny*, MetaDecoder, and SemiBin or the latter two, but without *binny* (_wo_bi) Upper panel: Reference genome fragmentation in number of contigs. Middle panel: Intersection size in number of NC MAGs with proportions of MAGs stemming from the six CAMI data sets. Lower panel: Number of MAGs per binning method on the left, intersections *>* 9 in the centre.

## Discussion

*binny* is a fully automated binning method, recovering unique information in form of complete and pure MAGs. It combines k-mer composition, read coverage and lineage-specific marker gene sets for iterative, non-linear dimension reduction of genomic signatures and subsequent automated contig clustering with cluster assessment. The low-dimensional embedding strategy to reduce large amounts of features has been used before for binning to aid the clustering of contigs [34, 59]. Clustering algorithms perform better in fewer dimensions, because distance information becomes increasingly imprecise at higher dimensions and the chance of random correlation between features rises [60].

While there are already binning methods available that make use of marker genes [14,38,61] and also lower dimensional embedding of contig features [61], *binny* uses a new and unique iterative embedding and clustering strategy. Importantly, it assesses clusters of contigs during its iterations, recognizing when further splitting of clusters is necessary. Of note, this lowers the complexity of each clustering task enabling *binny* to recover genomes that might not be separable with only a single embedding or clustering attempt. This seems to work particularly well for large, complex communities as shown with different CAMI data sets.

In combination with the ability (enabled by the marker gene approach) to incorporate also short informative contigs, which would be discarded by most other binning methods due to their applied contig length thresholds, *binny* is able to deal with highly fragmented genomes as shown for the CAMI samples. Of the tested binning methods, only CONCOCT was also able to deal with highly fragmented genomes. Although for the CAMI data sets, contigs below 1000 bp rarely made up >5% of the recovered MAGs size, *binny* assigned those usually with high precision (Supplementary Table 25). Additionally, *binny* performed also particularly well at recovering highly contiguous CAMI genomes. This can again be attributed to the ability to assess purity and completeness using the marker gene approach, here in particular for single-contig genomes.

*binny* also outperformed all other tested binning methods on the CAMI co-assemblies, where the added information provided by the coverage data from multiple samples substantially increased the overall performance. This is in line with previous studies observing additional discriminatory power of differential coverage depth compared with only sequence-based features [13,15]. Here again, *binny*’s iterative, supervised strategy seems well suited to the complexity of assemblies that contain highly fragmented genomes.

We also evaluated the effect of masking potentially disruptive sequence regions for the calculation of *k*-mer profiles. While the difference in performance with and without masking was not substantial, we believe that it reduces noise in the *k*-mer distributions of contigs from the same genome. One key reason for the small impact in the current setting might be the strong effect of the read coverage depth on the embedding and clustering procedure, which could outweigh the impact of the masked *k*-mer profile. Masking reads mapping to the disruptive regions, also modifying the depth information might increase its effectiveness and could be implemented in future versions.

It is generally advised [18,62] to make use of refinement methods, such as DAS tool and MetaWRAP here, which employ ensemble approaches to recover more complete and pure MAGs than the single binning methods alone. *binny* was shown to be an excellent addition to such approaches, because of its ability to recover large amounts of unique, pure and complete MAGs (Figures 3 and 5).

Finally, the results of the 105 metagenome benchmark show that *binny*’s performance translates to real-world scenarios, competing well with the latest methods on the recovery of NC and HQ MAG, while massively outperforming all other methods on the number of MIMAG-standard MAGs recovered. Still, there are also many samples where all binning methods were unable to recover a sizeable proportion of the assemblies as MAGs of sufficient quality. This might hint at the still limited capabilities of binning methods or could be caused by low quality of these assemblies.

## Conclusion

In conclusion, we demonstrate that *binny* outperforms or is highly competitive with currently available, state-of-the-art and/or popular binning methods based on established evaluation metrics, recovering unique, complete, and pure MAGs from simple and complex samples alike, while being able to handle contiguous, as well as fragmented genomes. Moreover, we could show that *binny* adds new MAGs when used in combination with other binning methods and binning refinement approaches, enabling researchers to further improve the recovery of genomes from their metagenomes.

---

**Additional file 1 — supplementary_figures.pdf**

Contains supplementary figures 1-23.

**Additional file 2 — supplementary_tables.xlsx**

Contains supplementary tables 1-25 as Excel sheets.

## Author’s contributions

O.H., P.M., and A.H.-B. designed this study. O.H. and A.H.-B. created the application. O.H. performed all experiments. O.H., P.M. and A.H.-B. wrote the manuscript; P.Q. and P.W. contributed to the review of the manuscript before submission. All authors read and approved the manuscript.

## Acknowledgements

The authors would like to thank Francesco Delogu and Benoit Kunath, for scientific discussions. Development and data analysis were performed using the research cluster of the Faculty of Science at the University of Amsterdam and the HPC facilities of the University of Luxembourg, both of whose administrators are thanked for excellent support. We also thank Adrian Fritz of the CAMI team for support.

## Availability of data and material

The latest version of *binny* can be found at https://github.com/a-h-b/binny. Scripts used in this study and related data is available at https://github.com/ohickl/binny_manuscript and https://doi.org/10.5281/zenodo.6977014.

## Footnotes

1 https://data.ace.uq.edu.au/public/CheckM_databases/

2 https://data.cami-challenge.org/participate

3 https://openstack.cebitec.uni-bielefeld.de:8080/swift/v1/CAMI_I_TOY_HIGH

4 https://gist.github.com/nathanhaigh/3521724#file-deinterleave_fastq-sh

5 https://jgi.doe.gov/

6 https://concoct.readthedocs.io/en/latest/usage.html

7 https://github.com/RasmussenLab/vamb/blob/master/README.md

9 https://github.com/pysam-developers/pysam

