## Supplementary figures 1-23 for "binny: an automated binning algorithm to recover high-quality genomes from complex metagenomic datasets"

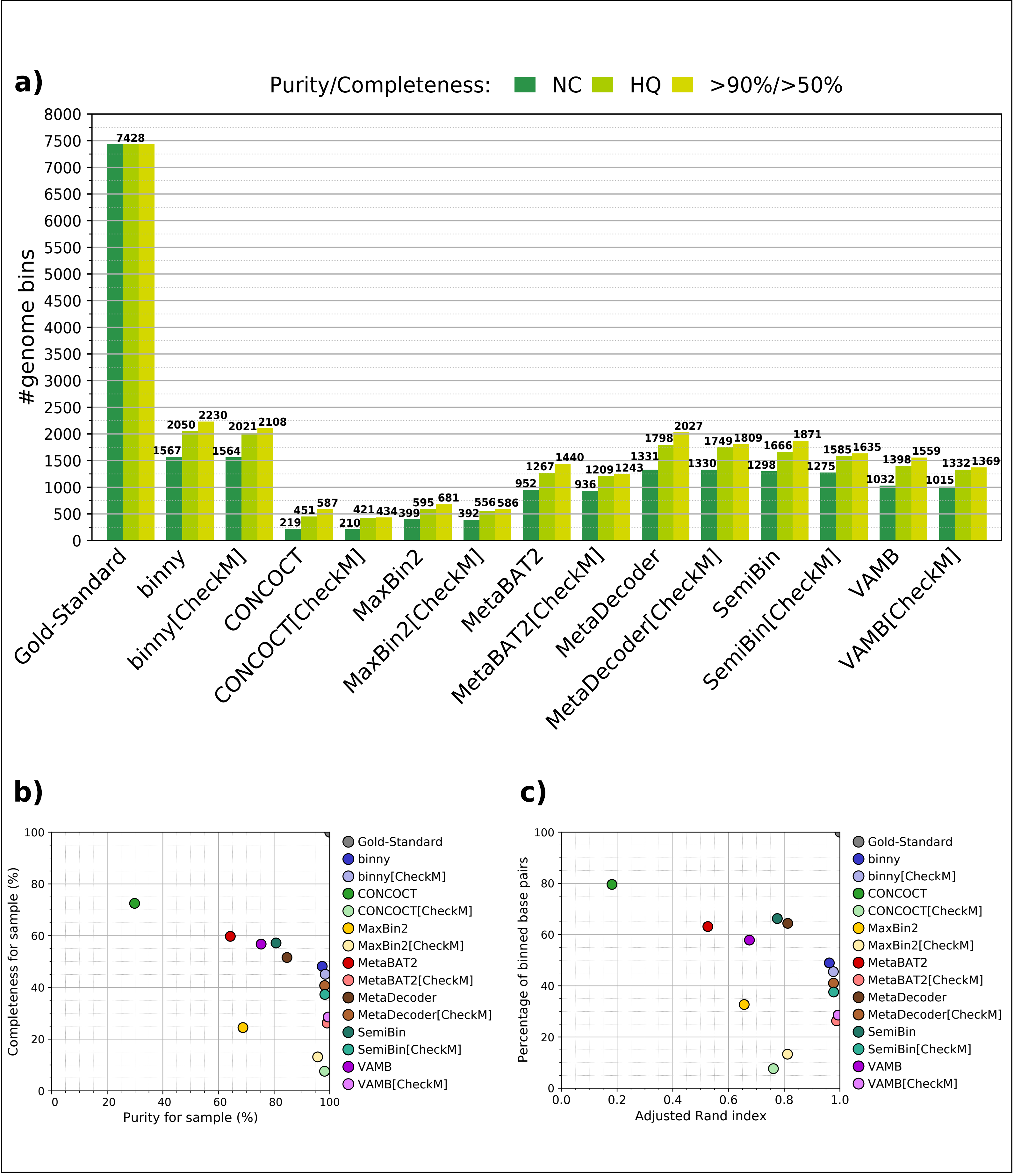


**Supplementary figure 1** AMBER binning stats for seven binning methods, with and without CheckM filtering, for 54 samples of six CAMI data sets. a) Numbers of MAGs recovered at near complete (NC, purity >95%, completeness >90%), high quality (HQ, purity >90%, completeness >70%), and purity >70%, completeness >50% quality thresholds. b) Average MAG completeness plotted against average purity. c) Average percentage of binned bp plotted against average Adjusted Rand Index.

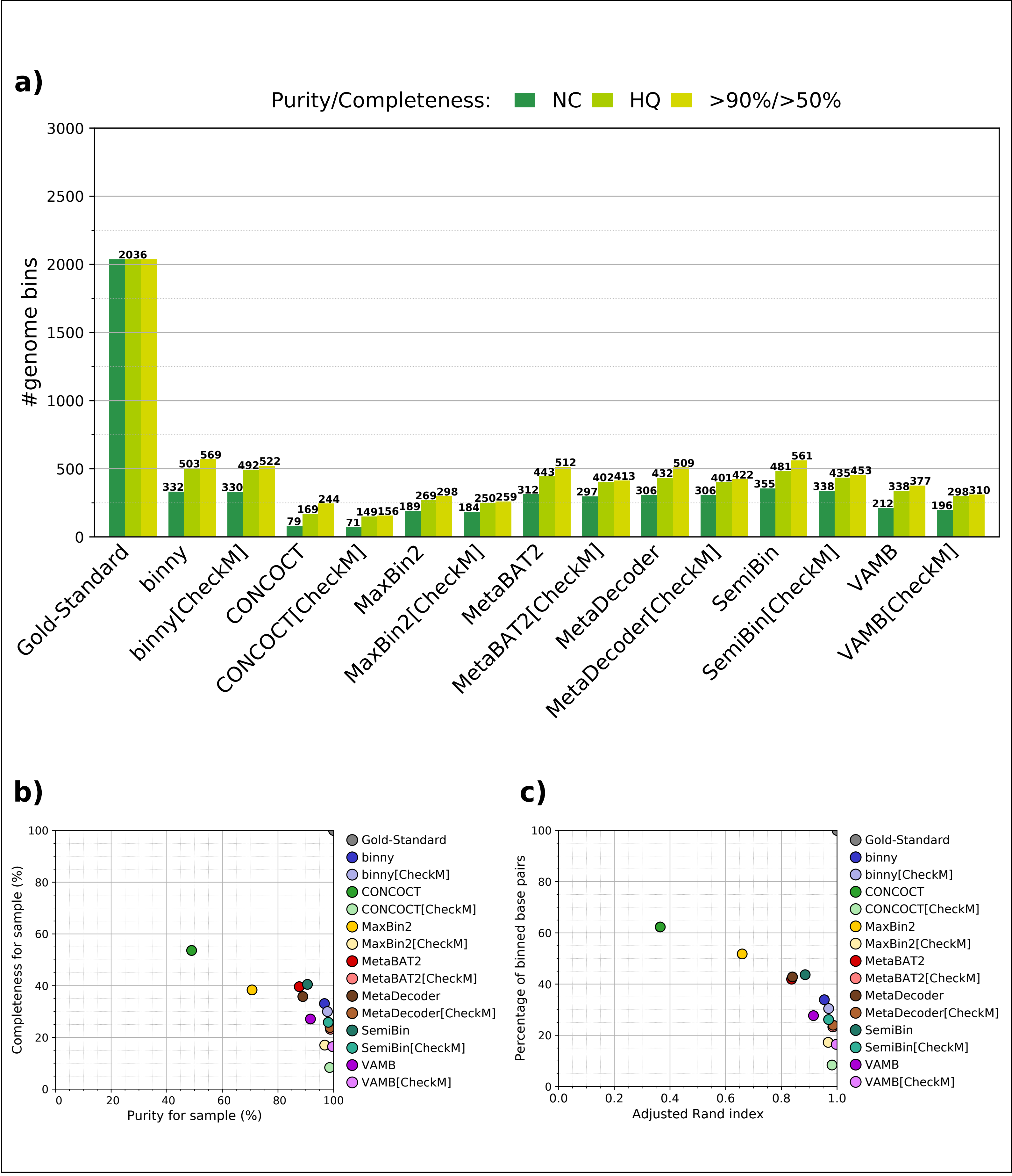


**Supplementary figure 2** AMBER binning stats for seven binning methods, with and without CheckM filtering, for the five samples of the CAMI 1 Toy High data set. a) Numbers of MAGs recovered at near complete (NC, purity >95%, completeness >90%), high quality (HQ, purity >90%, completeness >70%), and purity >70%, completeness >50% quality thresholds. b) Average MAG completeness plotted against average purity. c) Average percentage of binned bp plotted against average Adjusted Rand Index.

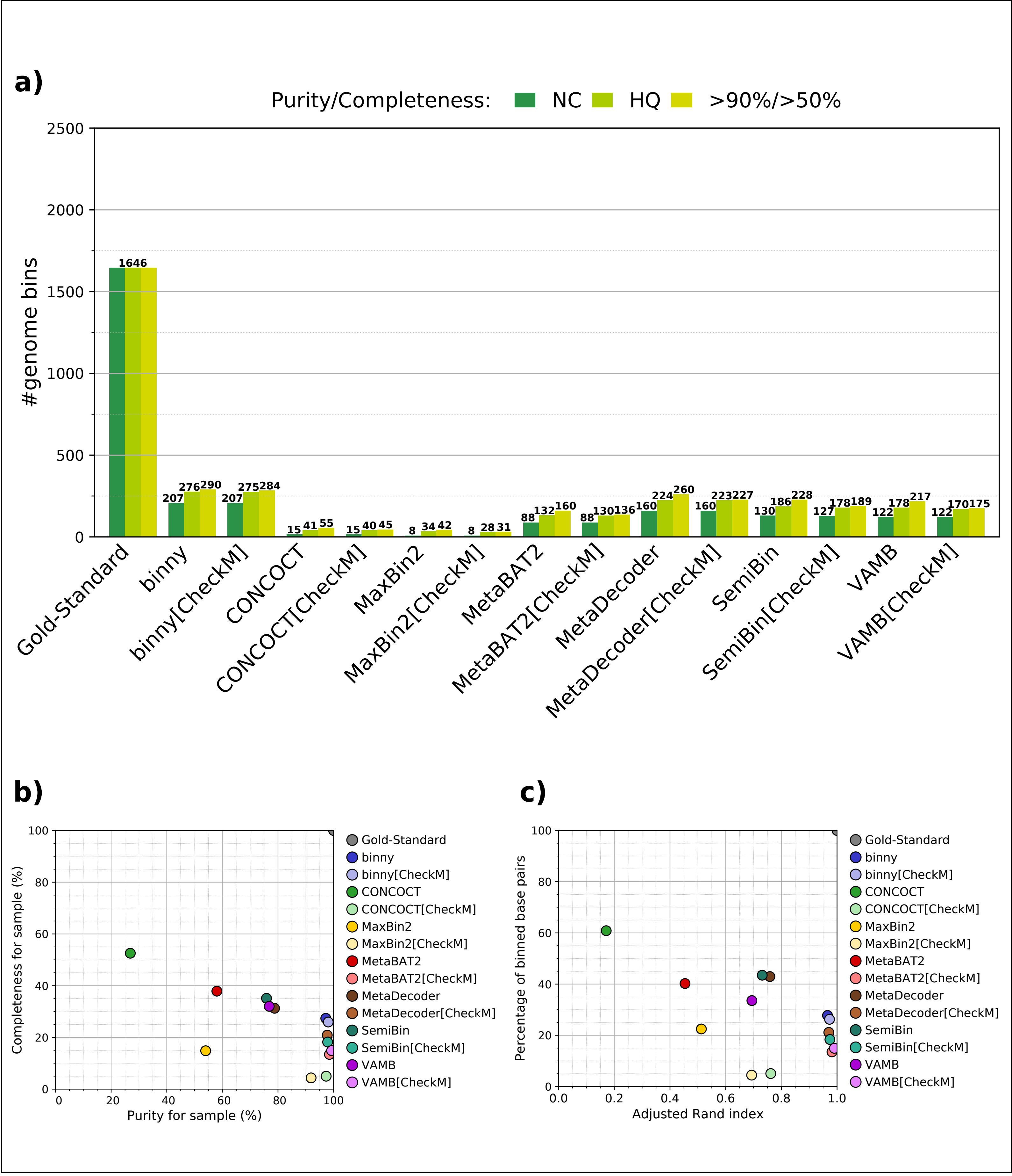


**Supplementary figure 3** AMBER binning stats for seven binning methods, with and without CheckM filtering, for the 10 samples of the CAMI 2 Toy Airways data set. a) Numbers of MAGs recovered at near complete (NC, purity >95%, completeness >90%), high quality (HQ, purity >90%, completeness >70%), and purity >70%, completeness >50% quality thresholds. b) Average MAG completeness plotted against average purity. c) Average percentage of binned bp plotted against average Adjusted Rand Index.

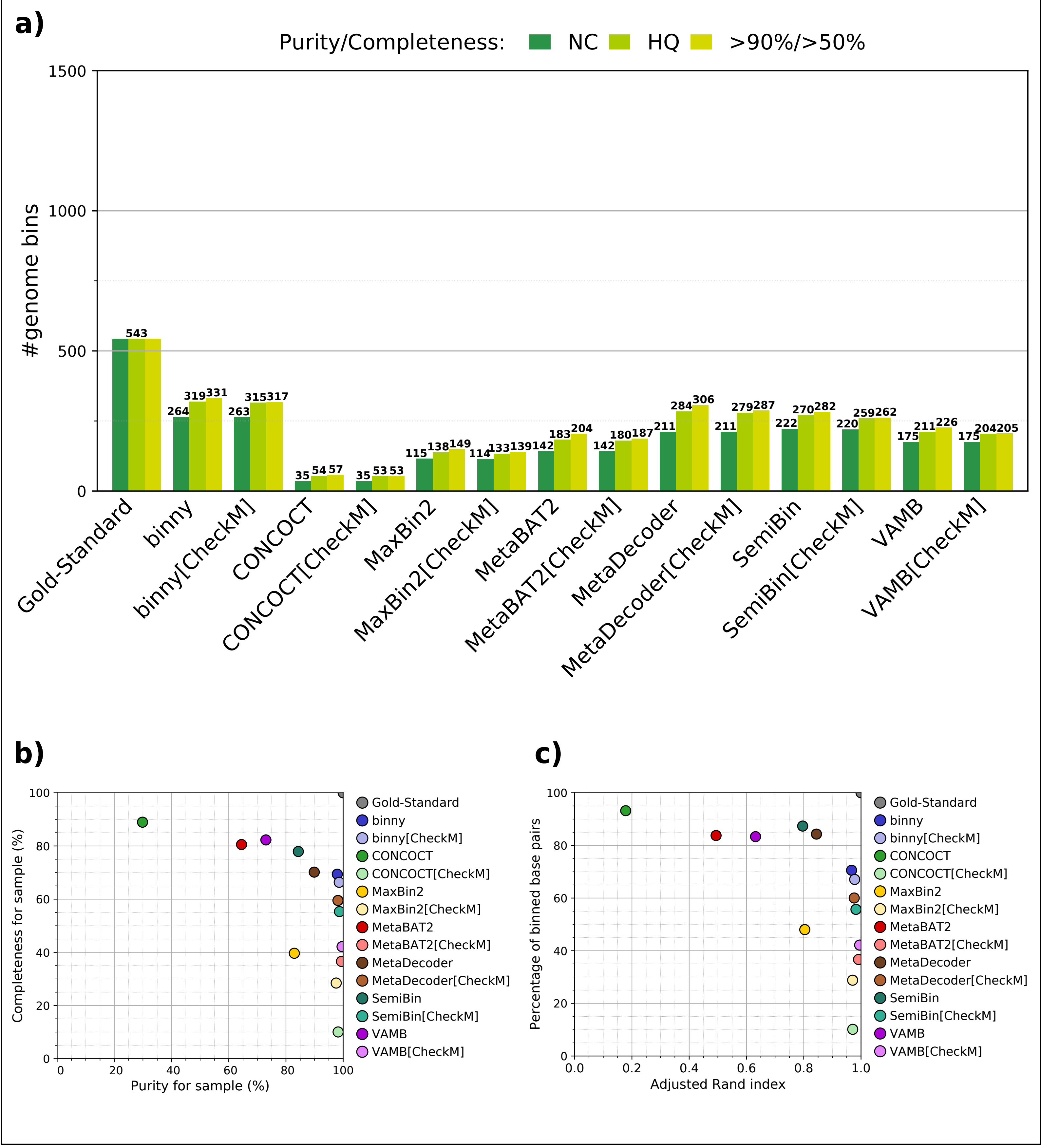


**Supplementary figure 4** AMBER binning stats for seven binning methods, with and without CheckM filtering, for the 10 samples of the CAMI 2 Toy Gastrointestinal data set. a) Numbers of MAGs recovered at near complete (NC, purity >95%, completeness >90%), high quality (HQ, purity >90%, completeness >70%), and purity >70%, completeness >50% quality thresholds. b) Average MAG completeness plotted against average purity. c) Average percentage of binned bp plotted against average Adjusted Rand Index.

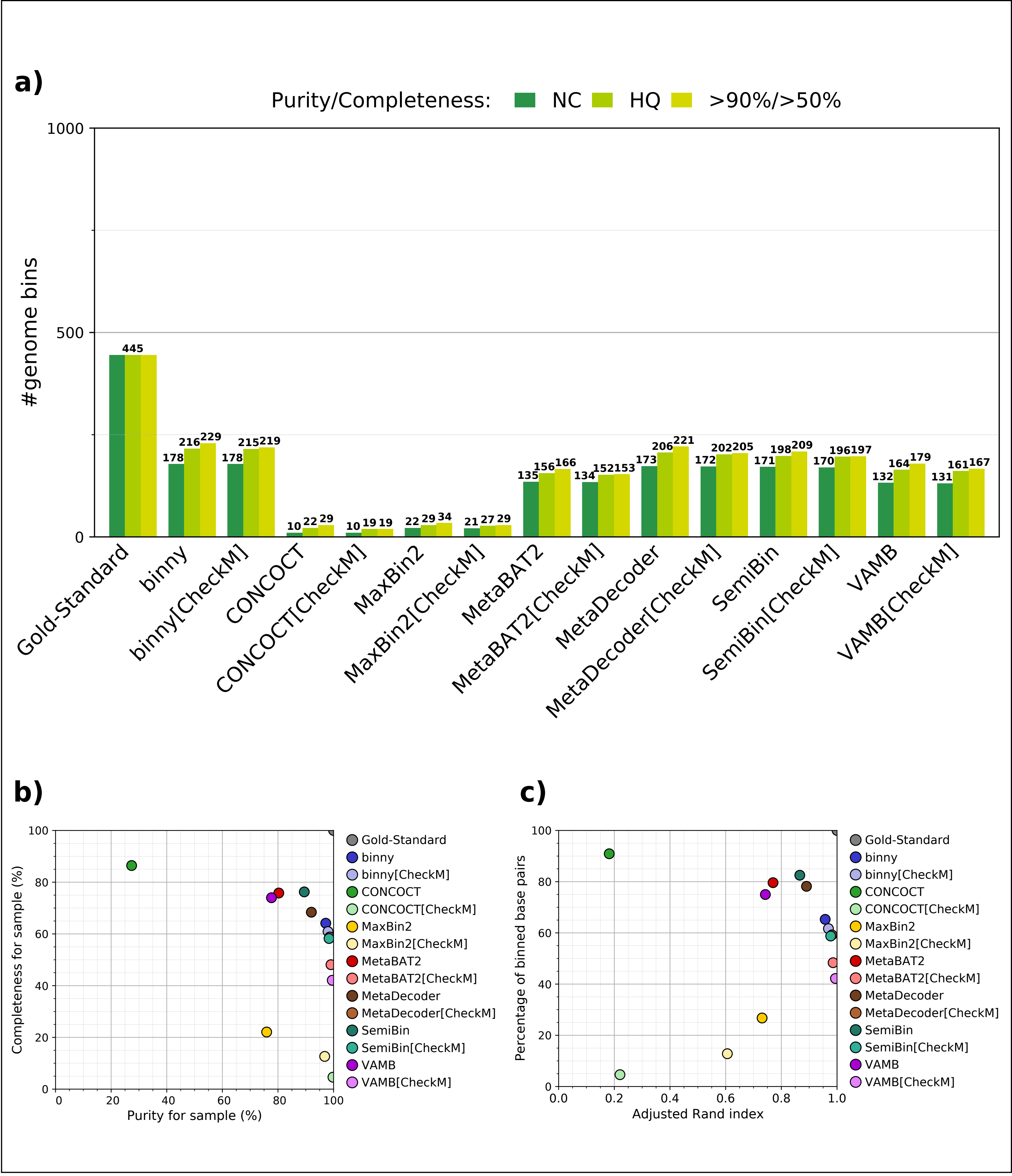


**Supplementary figure 5** AMBER binning stats for seven binning methods, with and without CheckM filtering, for the nine samples of the CAMI 2 Toy Urogential data set. a) Numbers of MAGs recovered at near complete (NC, purity >95%, completeness >90%), high quality (HQ, purity >90%, completeness >70%), and purity >70%, completeness >50% quality thresholds. b) Average MAG completeness plotted against average purity. c) Average percentage of binned bp plotted against average Adjusted Rand Index.

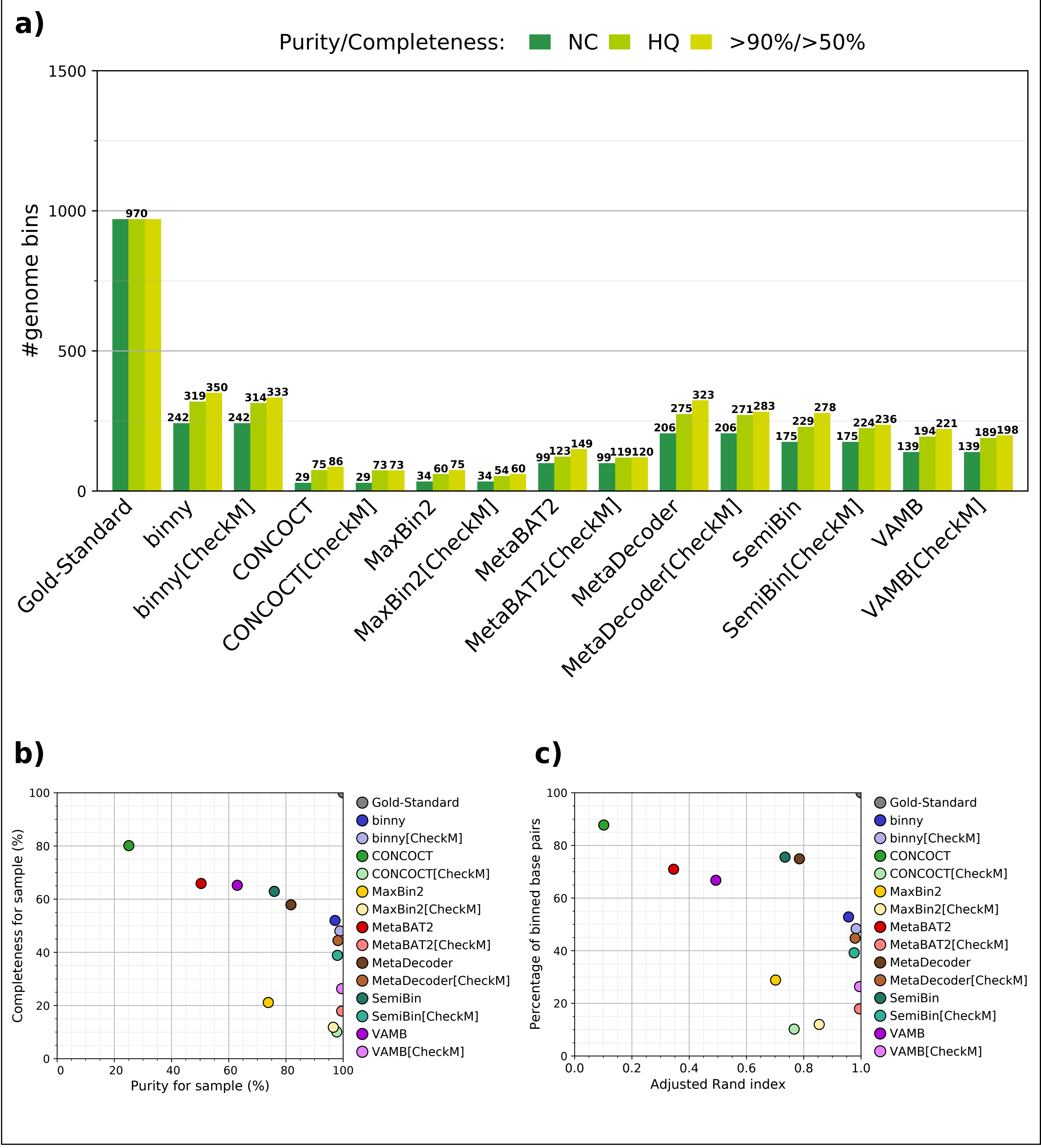


**Supplementary figure 6** AMBER binning stats for seven binning methods, with and without CheckM filtering, for the 10 samples of the CAMI 2 Toy Skin data set. a) Numbers of MAGs recovered at near complete (NC, purity >95%, completeness >90%), high quality (HQ, purity >90%, completeness >70%), and purity >70%, completeness >50% quality thresholds. b) Average MAG completeness plotted against average purity. c) Average percentage of binned bp plotted against average Adjusted Rand Index.

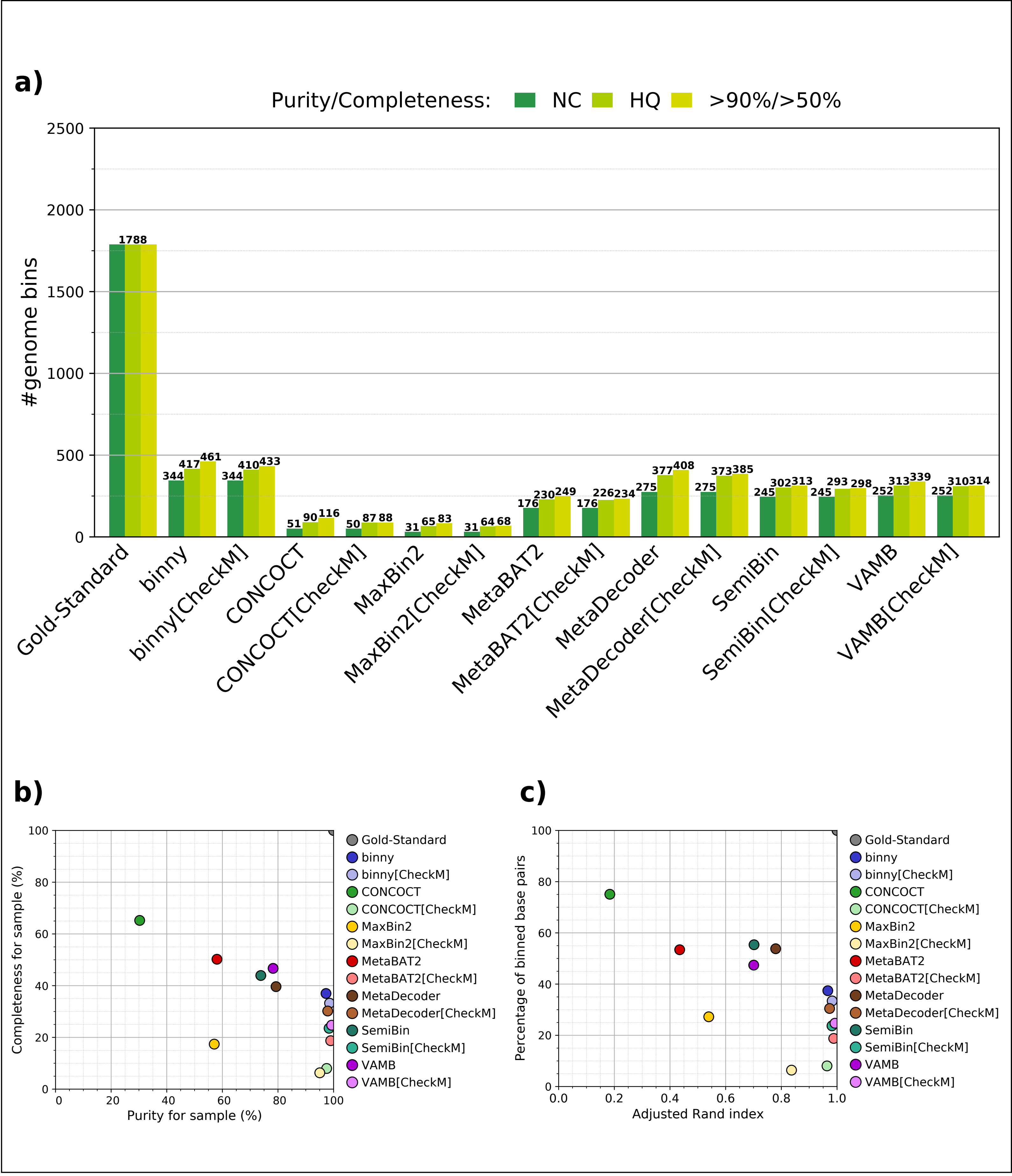


**Supplementary figure 7** AMBER binning stats for seven binning methods, with and without CheckM filtering, for the 10 samples of the CAMI 2 Toy Oral data set. a) Numbers of MAGs recovered at near complete (NC, purity >95%, completeness >90%), high quality (HQ, purity >90%, completeness >70%), and purity >70%, completeness >50% quality thresholds. b) Average MAG completeness plotted against average purity. c) Average percentage of binned bp plotted against average Adjusted Rand Index.

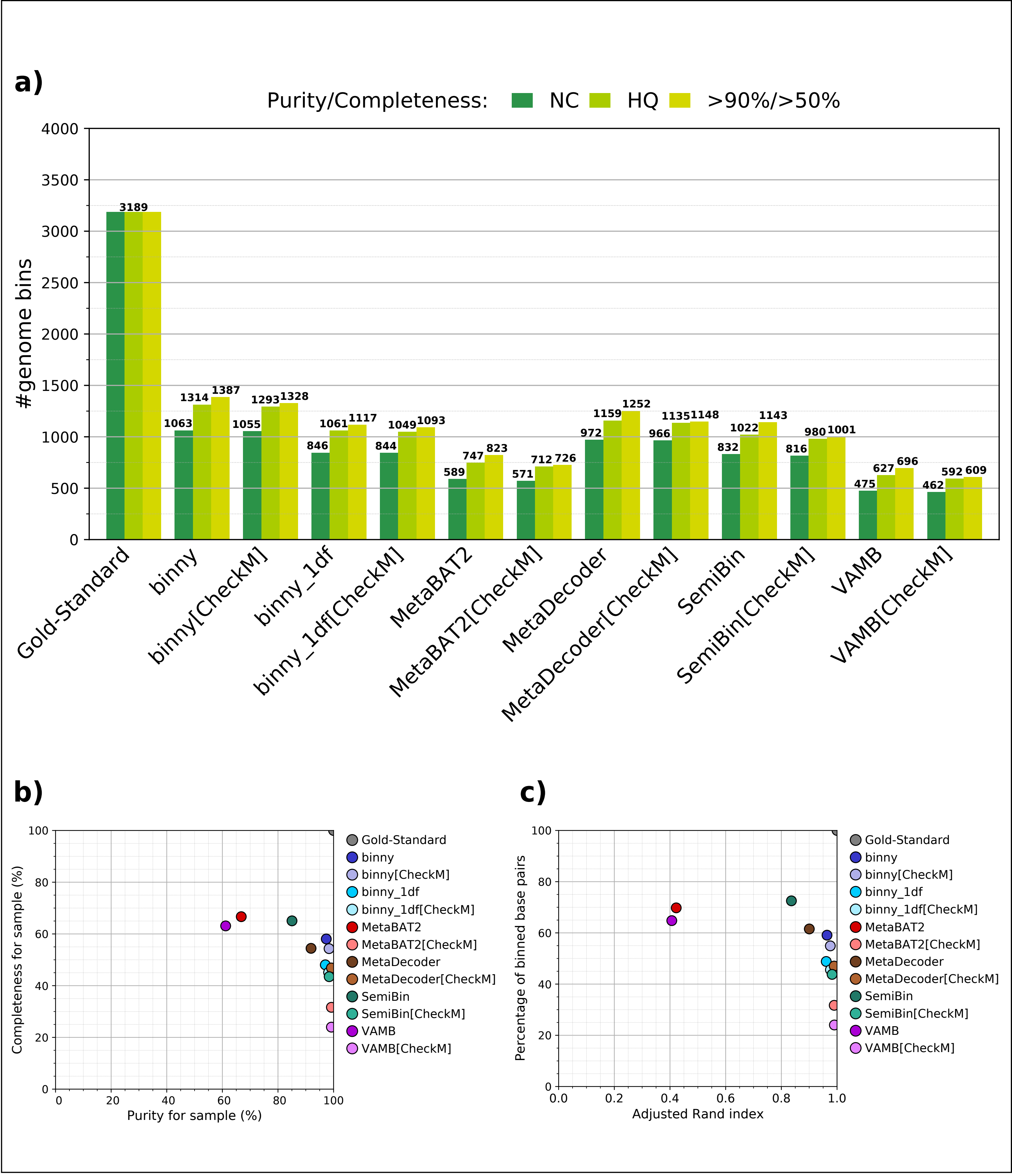


**Supplementary figure 8** AMBER binning stats for five binning methods and binny using only one depth file, with and without CheckM-filtering, for the six CAMI data set co-assemblies. a) Numbers of MAGs recovered at near complete (NC, purity >95%, completeness >90%), high quality (HQ, purity >90%, completeness >70%), and purity >70%, completeness >50% quality thresholds. b) Average MAG completeness plotted against average purity. c) Average percentage of binned bp plotted against average Adjusted Rand Index.

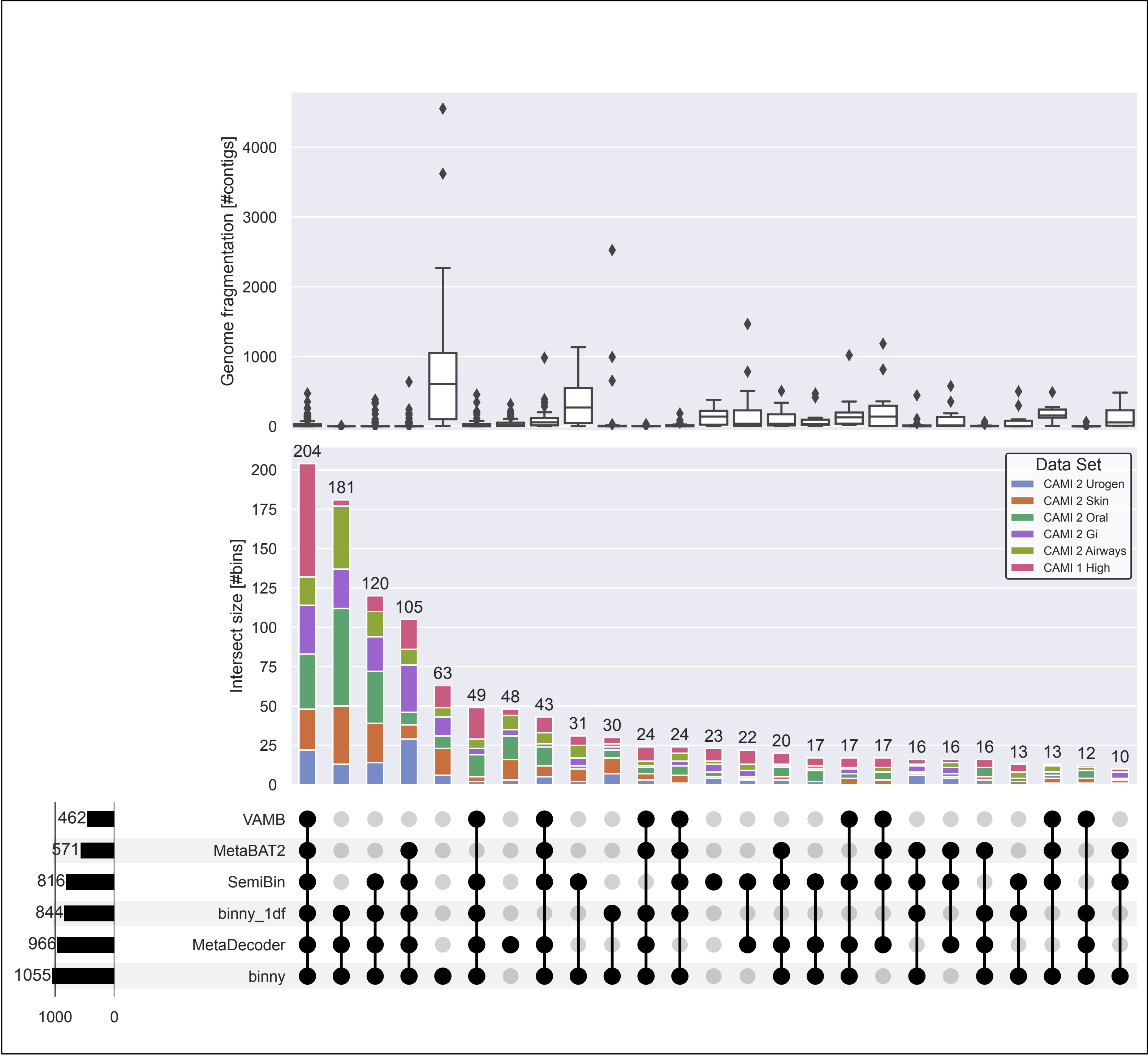


**Supplementary figure 9** UpSet plot of genome fragmentation and intersections of near complete (NC, purity >95%, completeness >90%) MAGs of five CheckM-filtered binning methods and binny using only one depth file for the six CAMI data set co-assemblies. Upper panel: Genome fragmentation in number of contigs per genome. Middle panel: Intersection size in number of NC MAGs with proportions of MAGs stemming from the six CAMI data sets. Lower panel: Number of MAGs per binning method on the left, intersection types > 9 in the centre.

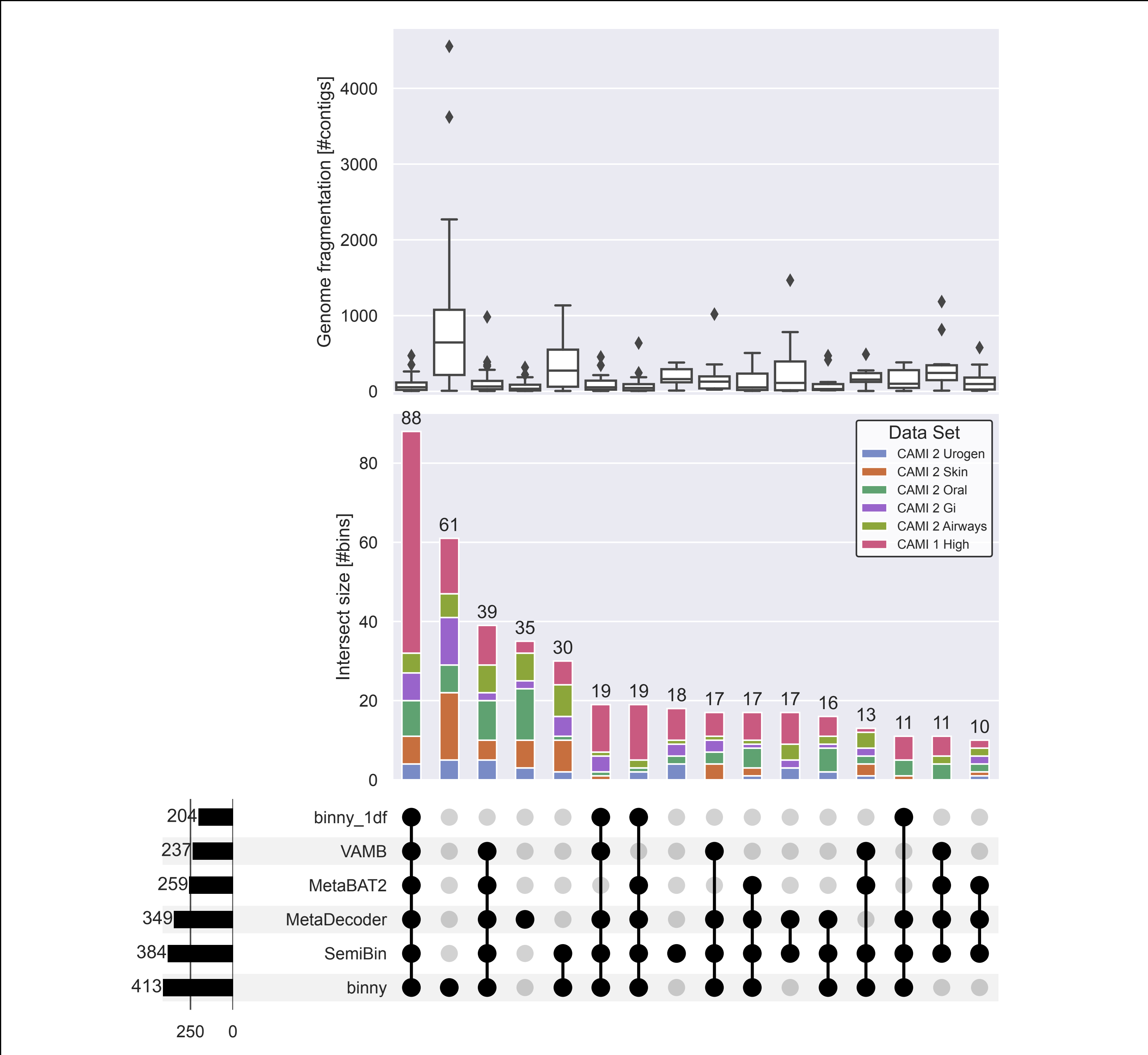


**Supplementary figure 10** UpSet plot of genome fragmentation and intersections of near complete (NC, purity >95%, completeness >90%) MAGs from genomes consisting of more than five contigs of five CheckM-filtered binning methods and binny using only one depth file for the six CAMI data set co-assemblies. Upper panel: Genome fragmentation in number of contigs per genome. Middle panel: Intersection size in number of NC MAGs with proportions of MAGs stemming from the six CAMI data sets. Lower panel: Number of MAGs per binning method on the left, intersection types > 9 in the centre.

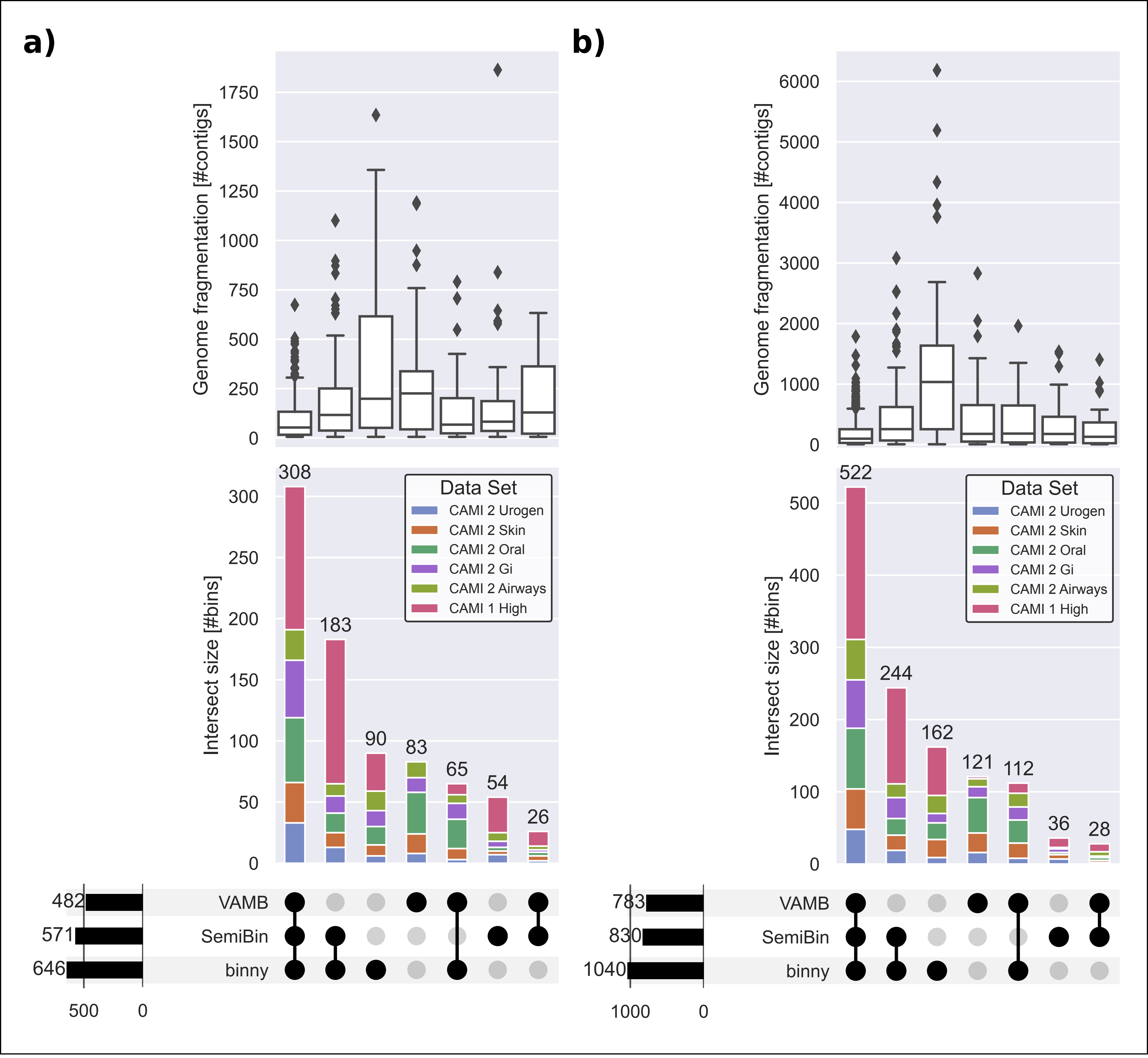


**Supplementary figure 11** UpSet plots of genome fragmentation and intersections of a) near complete (NC, purity>95%, completeness >90%), and b) high quality (HQ, purity >90%, completeness >70%) MAGs from genomes consisting of more than five contigs of three CheckM-filtered binning methods for 54 samples from six CAMI data sets. Upper panels: Genome fragmentation in number of contigs per genome. Middle panels: Intersection size in number of a) NC, b) HQ MAGs with proportions of MAGs stemming from the six CAMI data sets. Lower panels: Number of MAGs per binning method on the left, intersection types > 9 in the centre.

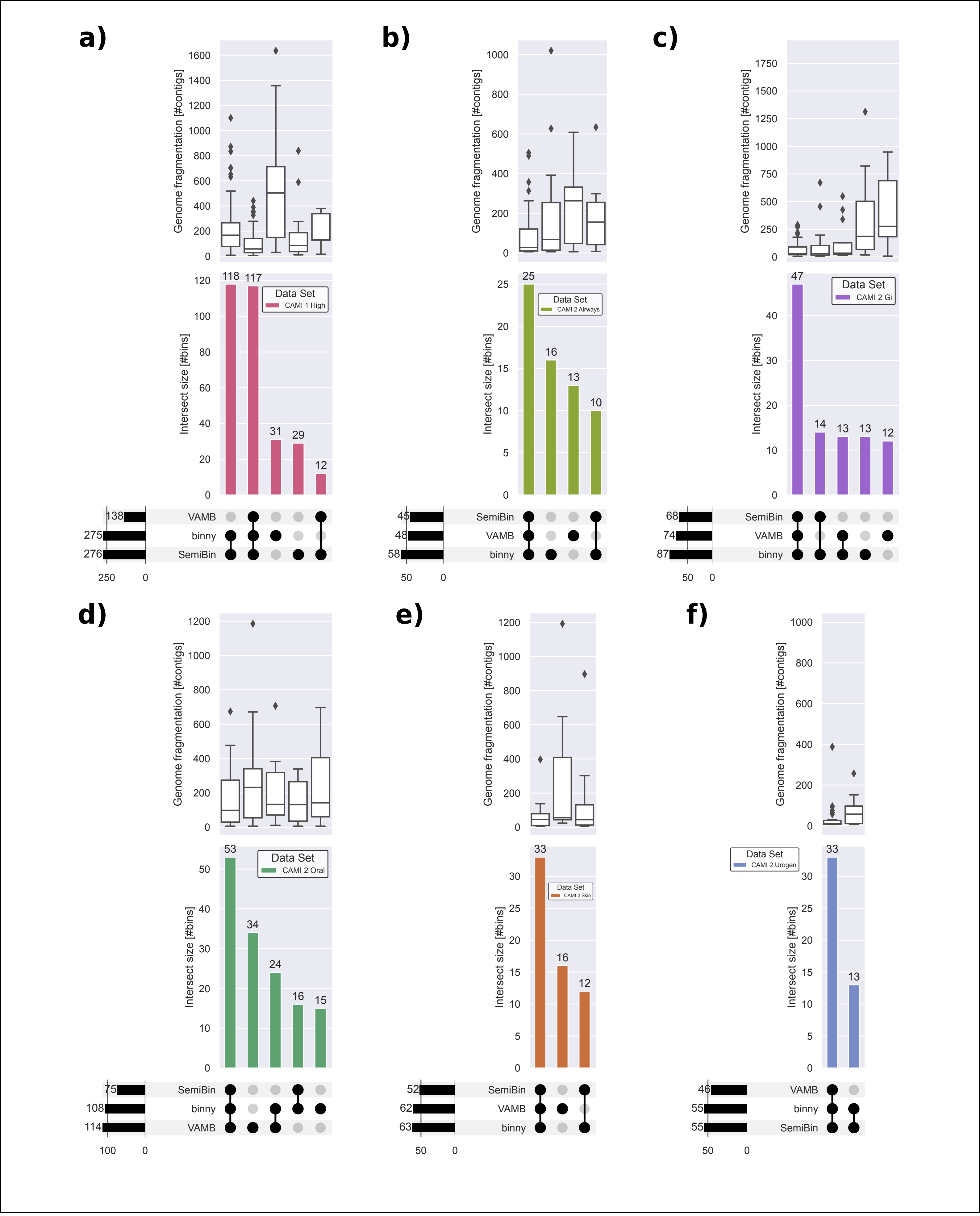


**Supplementary figure 12** UpSet plots of genome fragmentation and intersections of near complete (NC, purity >95%, completeness >90%) MAGs from genomes consisting of more than five contigs of three CheckM-filtered binning methods for 54 samples from a) CAMI 1 Toy High, b) CAMI 2 Toy Airways, c) CAMI 2 Toy Gastrointestinal, d) CAMI 2 Toy Oral, e) CAMI 2 Toy Skin, and f) CAMI 2 Toy Urogen data sets. Upper panels: Genome fragmentation in number of contigs per genome. Middle panels: Intersection size in number of NC MAGs with proportions of MAGs stemming from the six CAMI data sets. Lower panels: Number of MAGs per binning method on the left, intersection types > 9 in the centre.

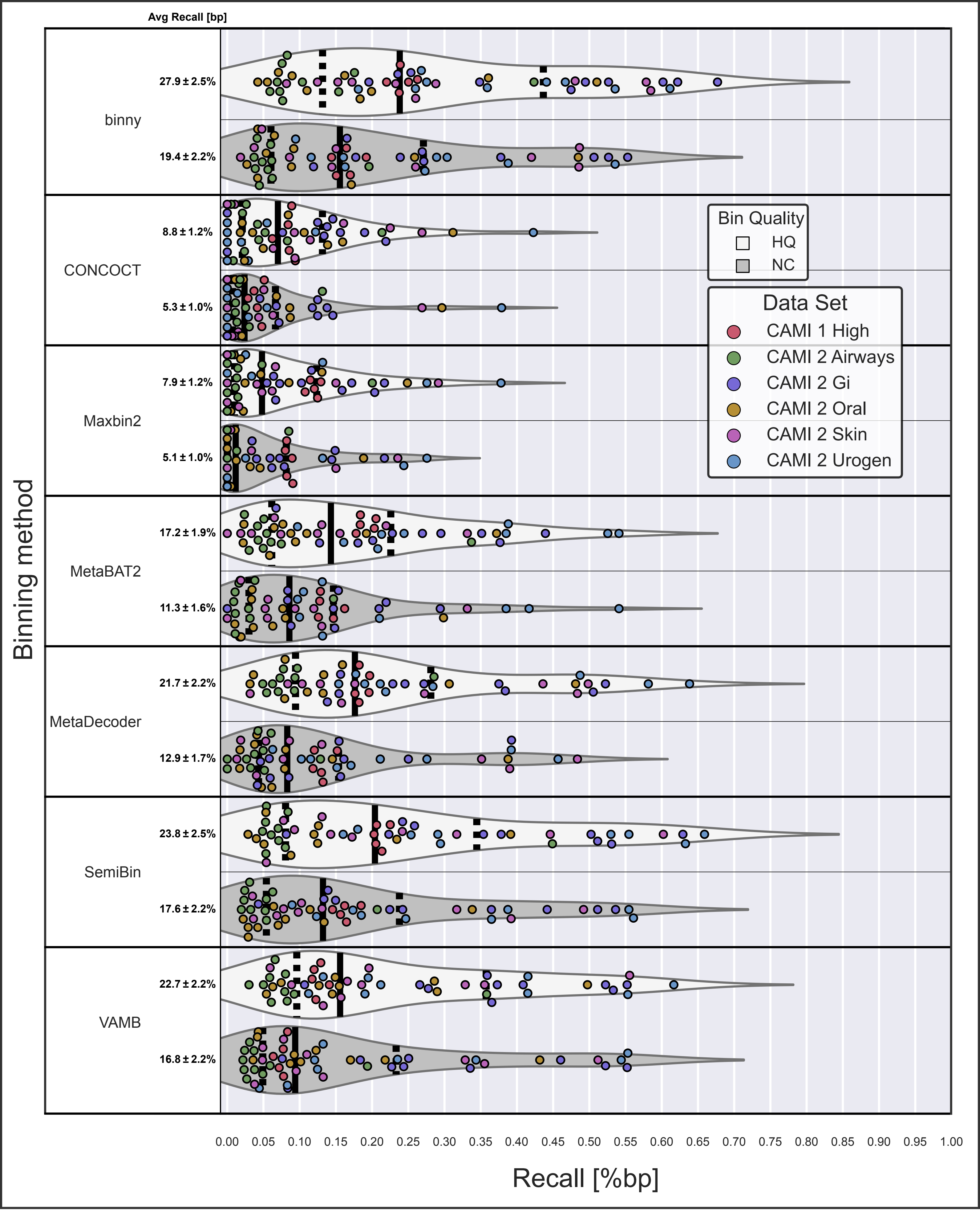


**Supplementary figure 13** Recall of bp assembled sequences as HQ (high-quality, purity >90%, completeness >70%) and NC (near-complete, purity >95%, completeness >90%) MAGs of genomes consisting of more than five contigs per binning method per sample, for the six CAMI data sets. The average recall is shown with the standard errors of the mean.

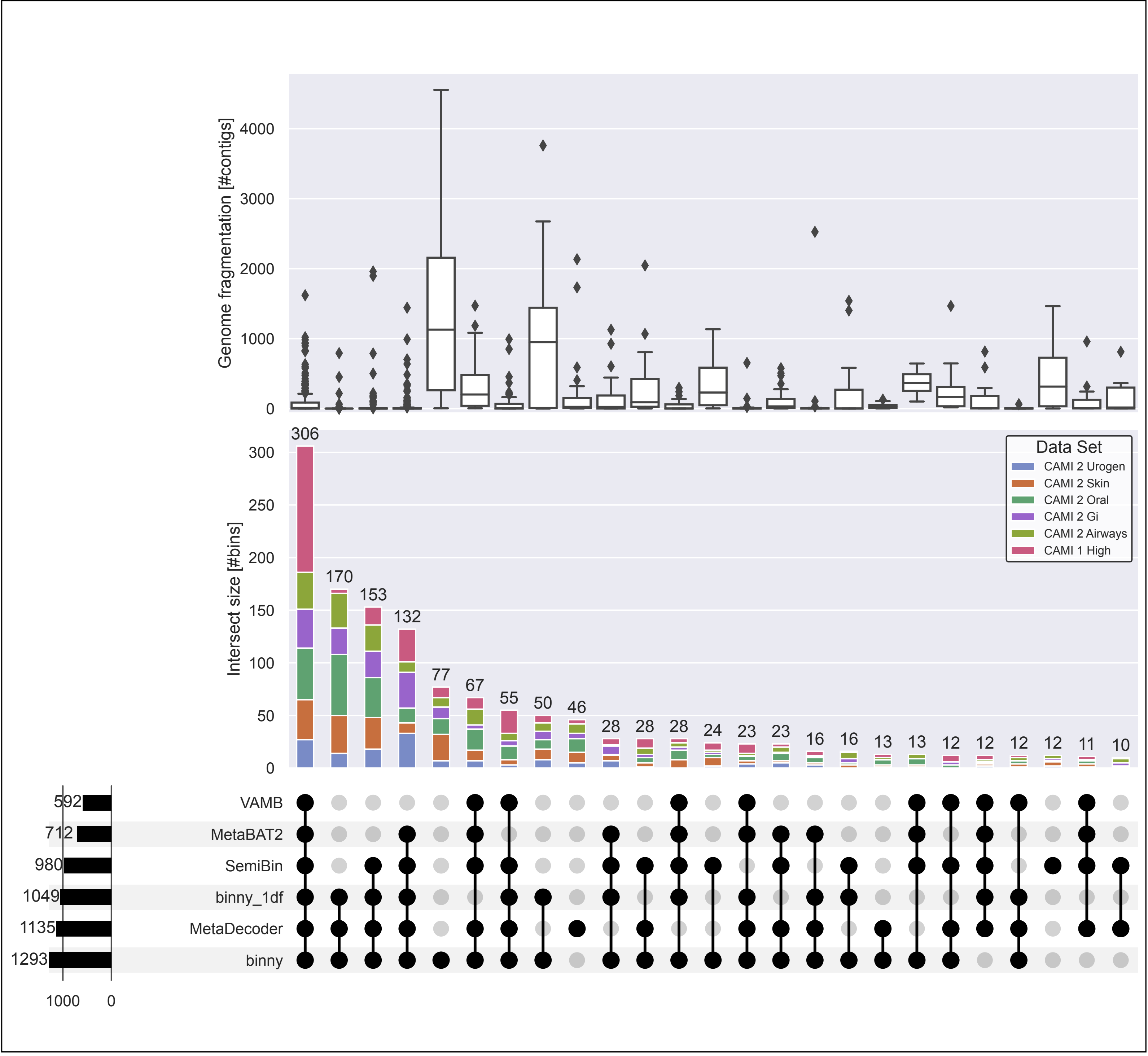


**Supplementary figure 14** UpSet plot of genome fragmentation and intersections of high quality (HQ, purity >90%, completeness >70%) bins of five CheckM-filtered binning methods and binny using only one depth file for the six CAMI data set co-assemblies. Upper panel: Genome fragmentation in number of contigs per genome. Middle panel: Intersection size in number of HQ MAGs with proportions of MAGs stemming from the six CAMI data sets. Lower panel: Number of MAGs per binning method on the left, intersection types > 9 in the centre.

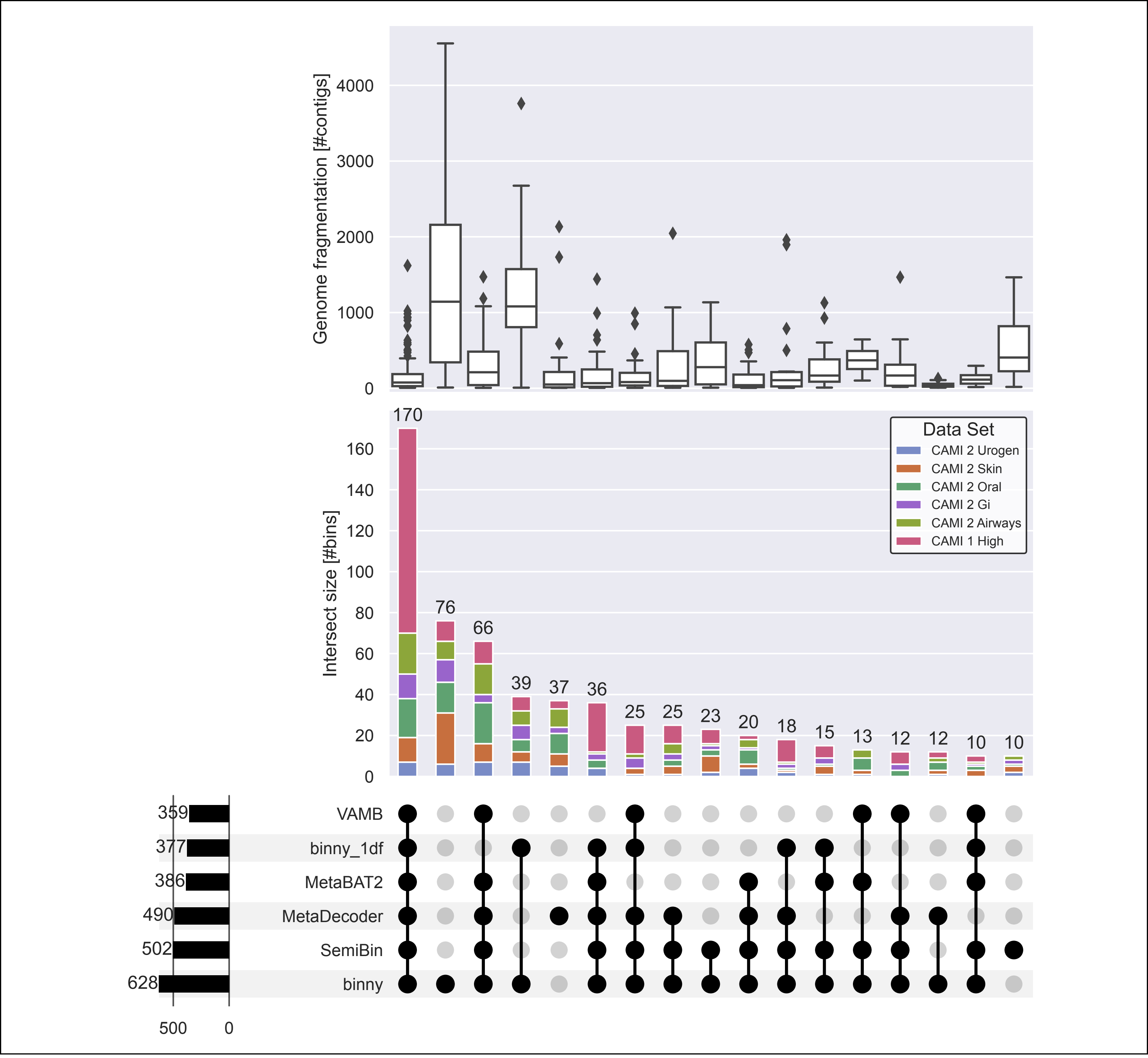


**Supplementary figure 15** UpSet plot of genome fragmentation and intersections of high quality (HQ, purity >90%, completeness >70%) MAGs of genomes consisting of more than five contigs of five CheckM-filtered binning methods and binny using only one depth file for six CAMI co-assemblies. Upper panel: Genome fragmentation in number of contigs per genome. Middle panel: Intersection size in number of HQ MAGs with proportions of MAGs stemming from the six CAMI data sets. Lower panel: Number of MAGs per binning method on the left, intersection types > 9 in the centre.

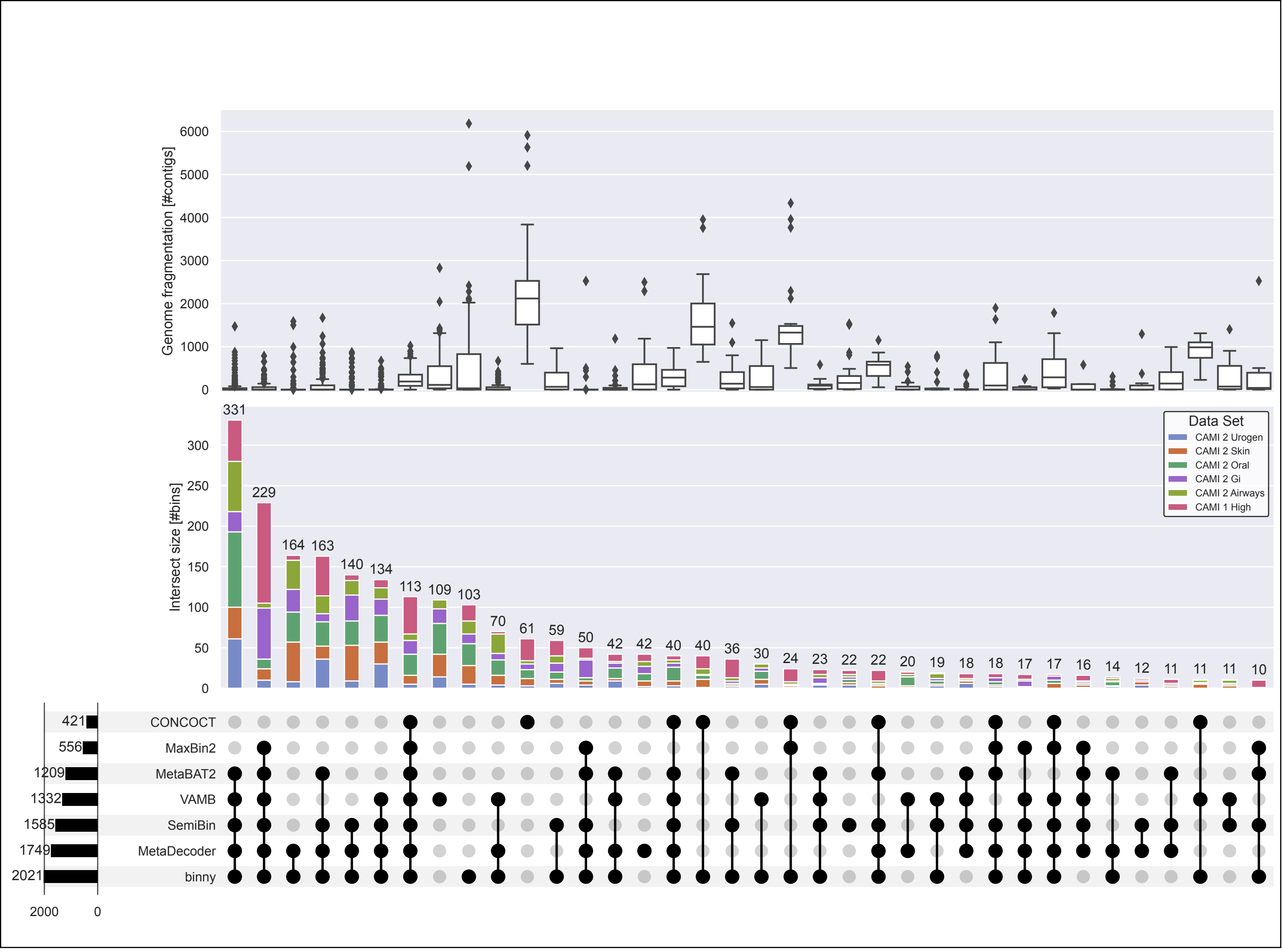


**Supplementary figure 16** UpSet plot of genome fragmentation and intersections of high quality (HQ, purity >90%, completeness >70%) MAGs of seven CheckM-filtered binning methods for 54 samples of six CAMI data sets. Upper panel: Genome fragmentation in number of contigs per genome. Middle panel: Intersection size in number of HQ MAGs with proportions of MAGs stemming from the six CAMI data sets. Lower panel: Number of MAGs per binning method on the left, intersection types > 9 in the centre.

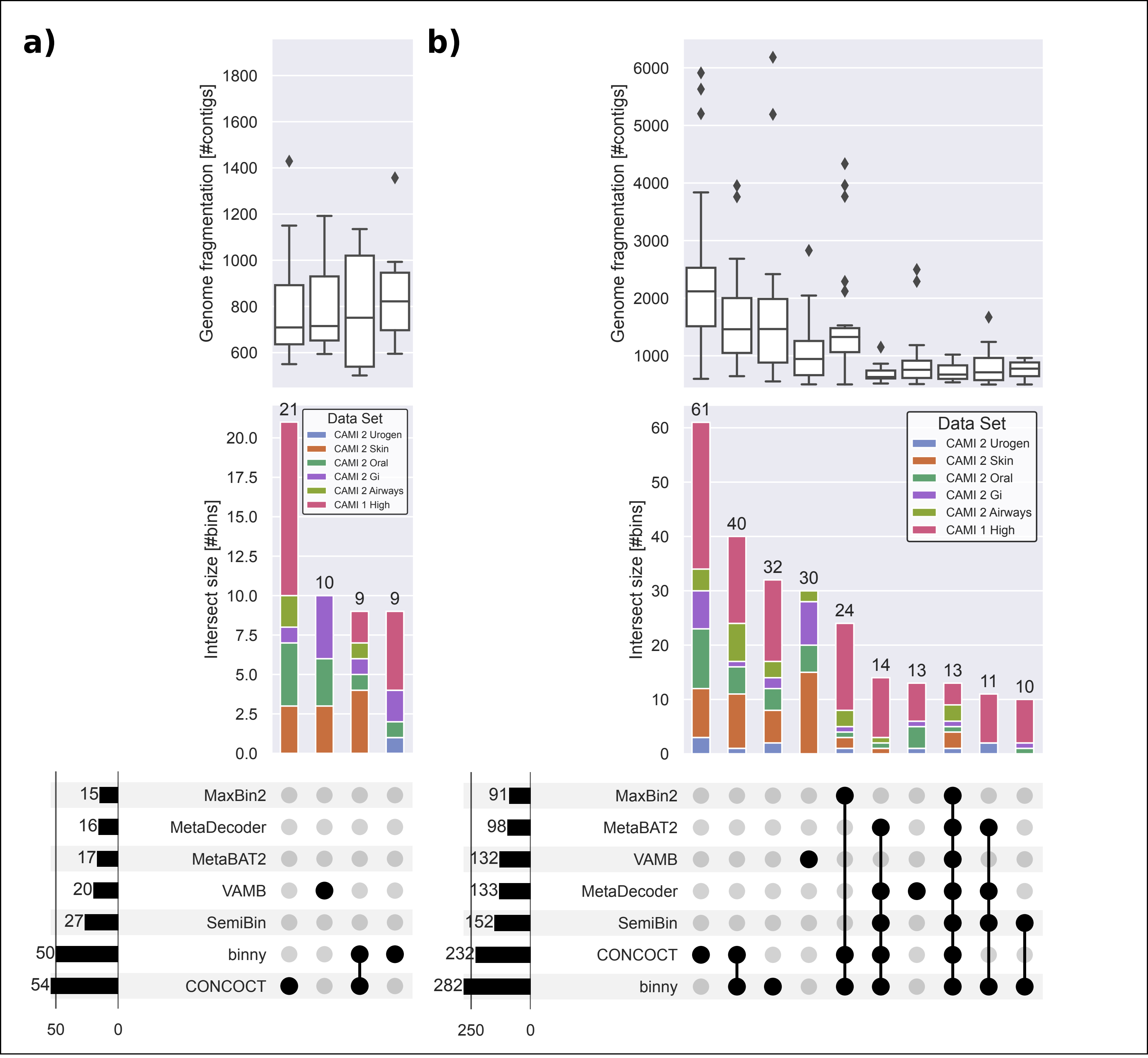


**Supplementary figure 17** UpSet plots of genome fragmentation and intersections of a) near complete (NC, purity >95%, completeness >90%), and b) high quality (HQ, purity >90%, completeness >70%) MAGs of genomes consisting of more than 500 contigs of seven CheckM-filtered binning methods for 54 samples from six CAMI data sets. Upper panels: Genome fragmentation in number of contigs per genome. Middle panels: Intersection size in number of a) NC, b) HQ MAGs with proportions of MAGs stemming from the six CAMI data sets. Lower panels: Number of MAGs per binning method on the left, intersection types a) > 5, b) > 9 in the centre.

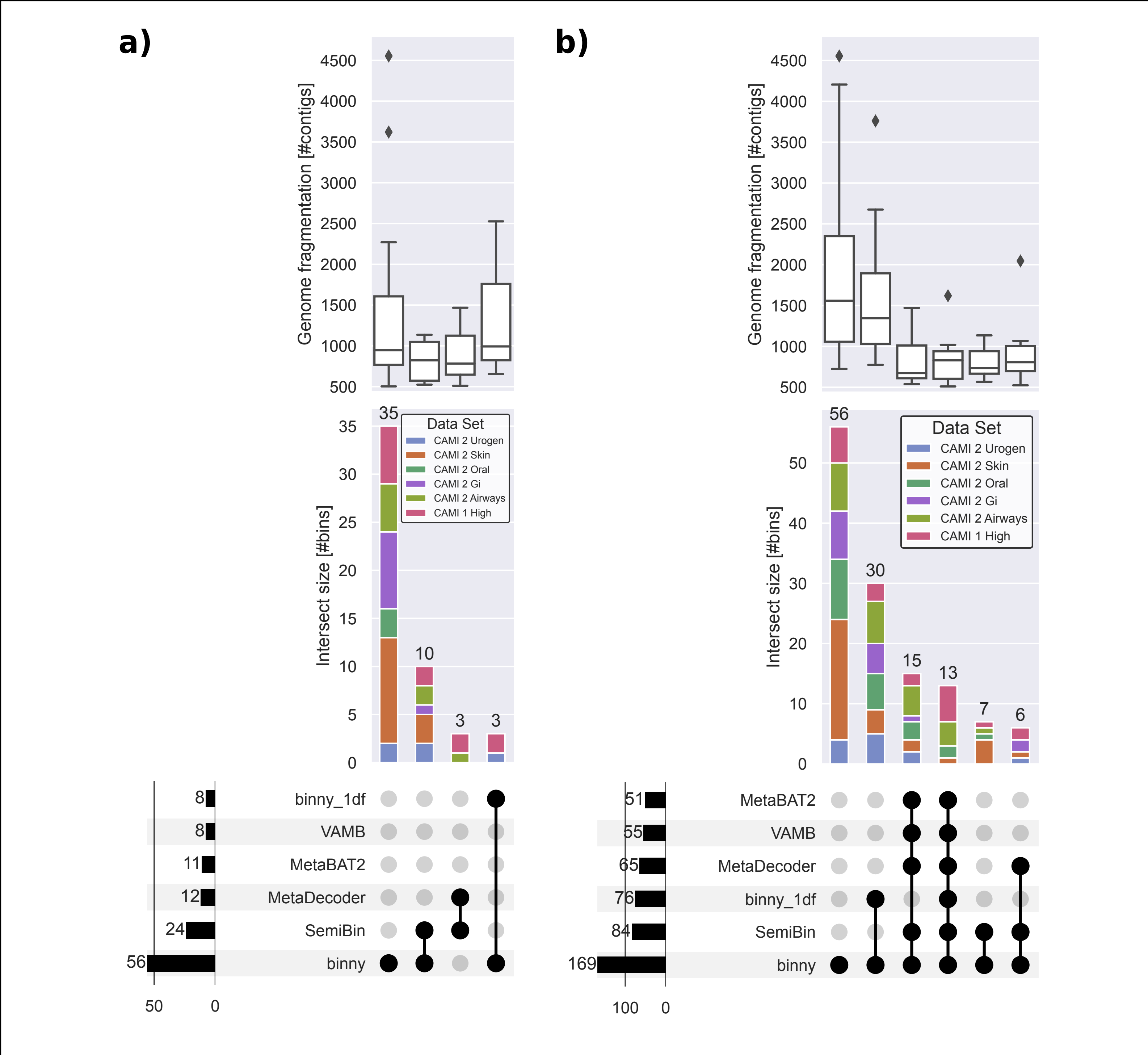


**Supplementary figure 18** UpSet plots of genome fragmentation and intersections of a) near complete (NC, purity >95%, completeness >90%), and b) high quality (HQ, purity >90%, completeness >70%) MAGs of genomes consisting of more than 500 contigs of five CheckM-filtered binning methods and binny using only one depth file for the six CAMI data set co-assemblies. Upper panels: Genome fragmentation in number of contigs per genome. Middle panels: Intersection size in number of a) NC, b) HQ MAGs with proportions of MAGs stemming from the six CAMI data sets. Lower panels: Number of MAGs per binning method on the left, intersection types a) > 2, b) > 4 in the centre.

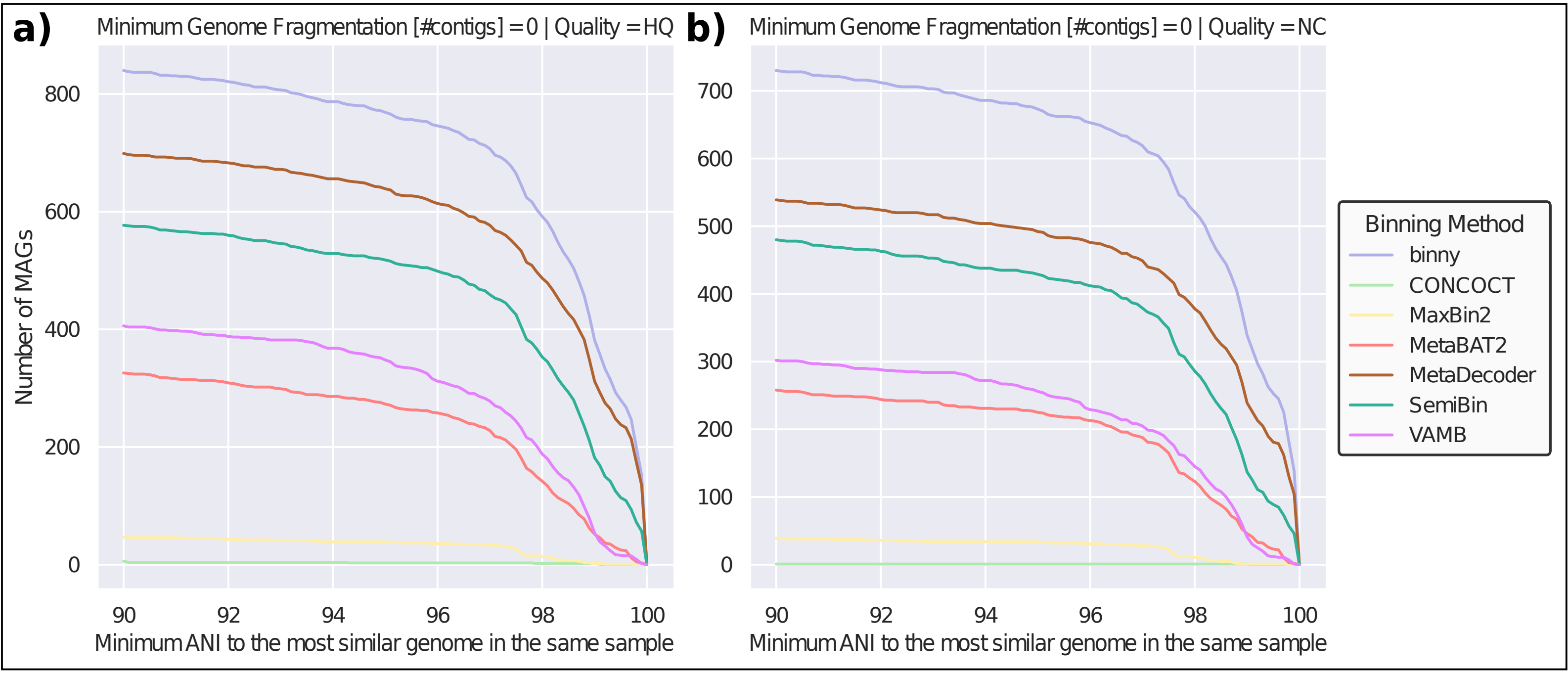


**Supplementary figure 19** Number of a) high quality (HQ, purity >90%, completeness >70%), and b) near complete (NC, purity >95%, completeness >90%) MAGs with a minimum ANI to the most similar genome in the same CAMI sample of at least 90.0% up to 99.9% in 0.1% steps for seven CheckM-filtered binning methods.

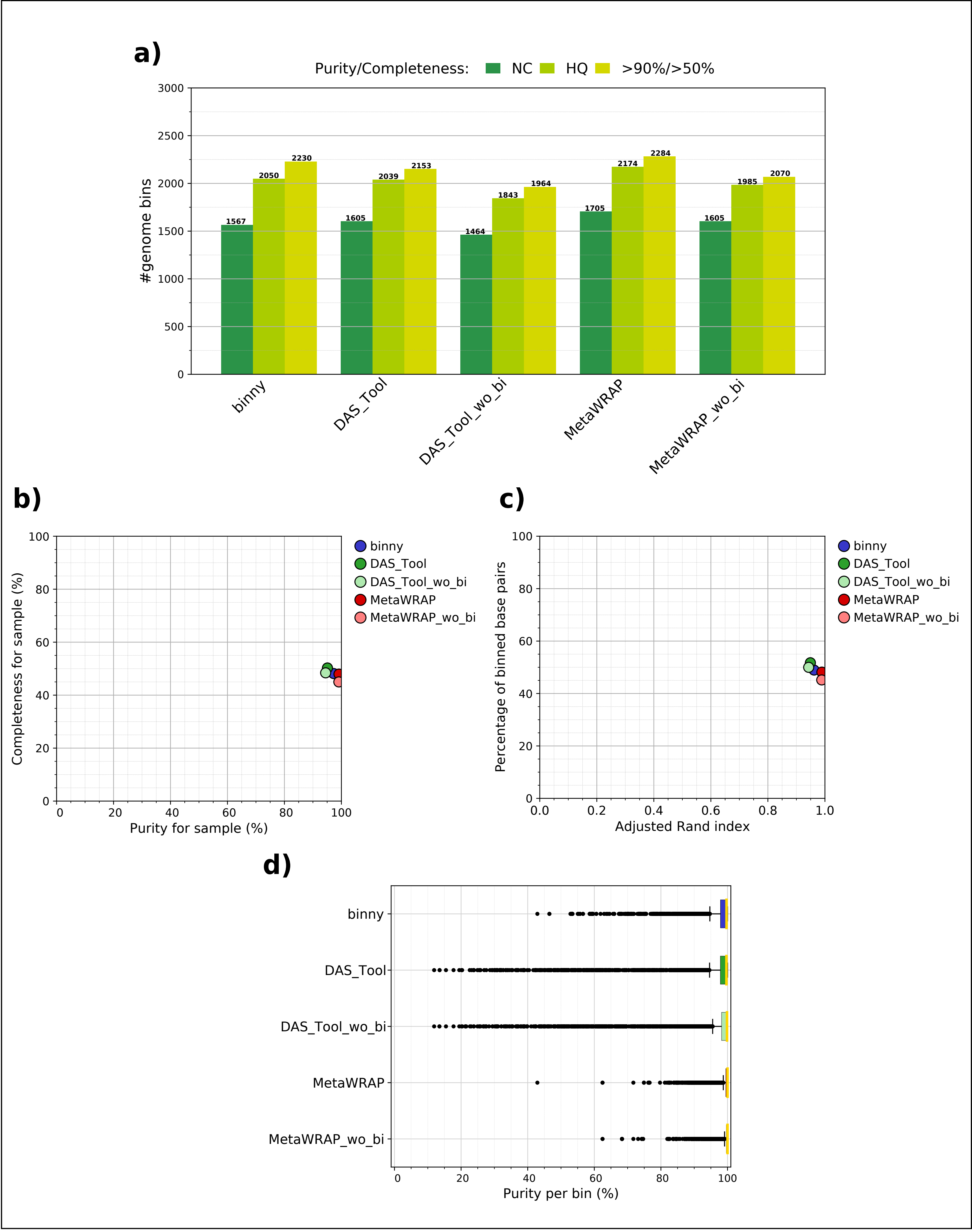


**Supplementary figure 20** AMBER binning stats for unfiltered binny and two binning refinement methods for 54 samples of six CAMI data sets. a) Numbers of MAGs recovered at near complete (NC, purity >95%, completeness >90%), high quality (HQ, purity >90%, completeness >70%), and purity >70%, completeness >50% quality thresholds. b) Average MAG completeness plotted against average purity c) Average percentage of binned bp plotted against average Adjusted Rand Index. d) Boxplot of MAG purity per method.

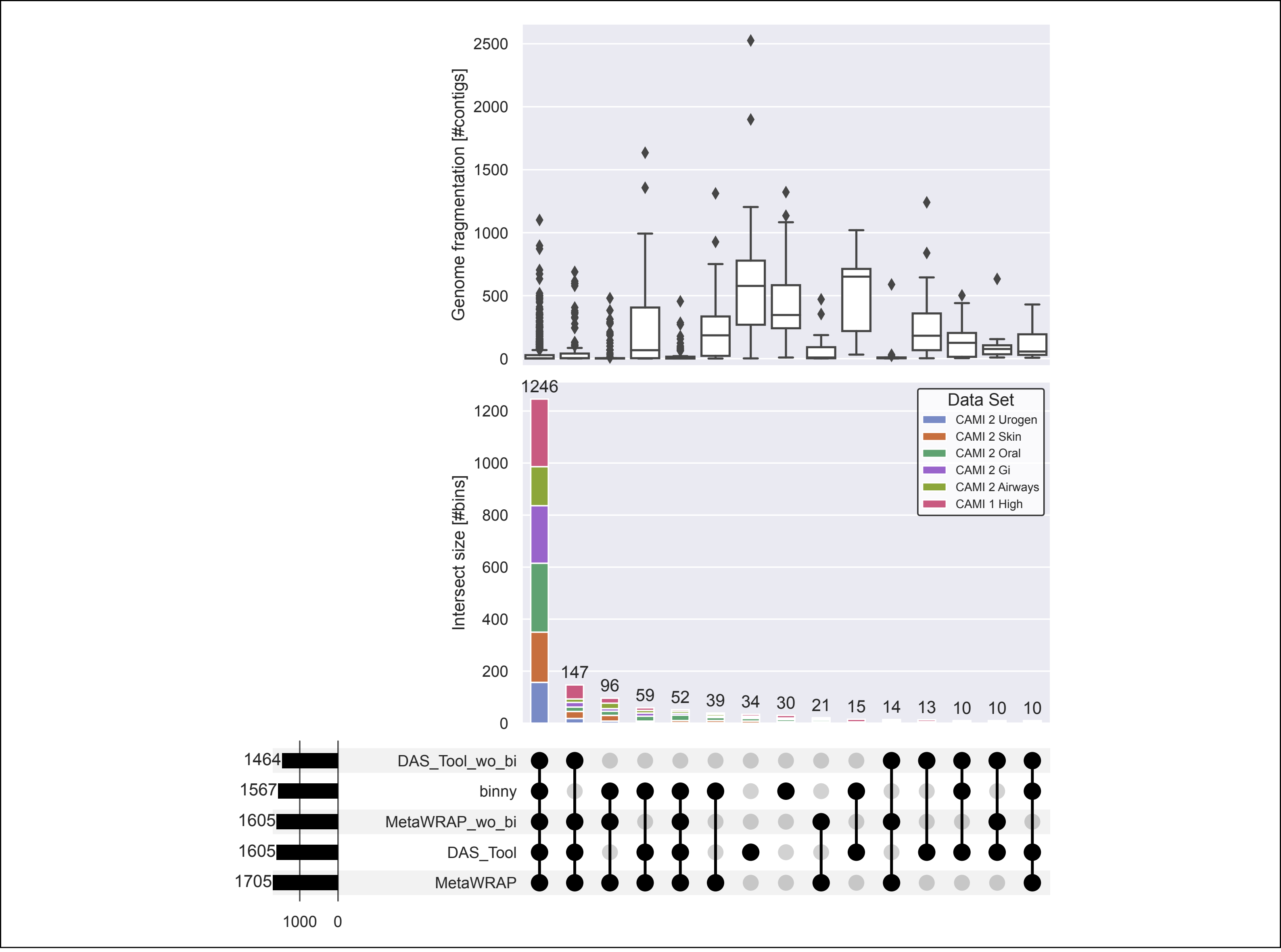


**Supplementary figure 21** UpSet plot of genome fragmentation and intersections of near complete (NC, purity >95%, completeness >90%) MAGs of unfiltered binny and two binning refinement methods for 54 samples from six CAMI data sets. Upper panel: Genome fragmentation in number of contigs. Middle panel: Intersection size in number of NC MAGs with proportions of MAGs stemming from the six CAMI data sets. Lower panel: Number of MAGs per binning method on the left, intersection types > 9 in the centre.

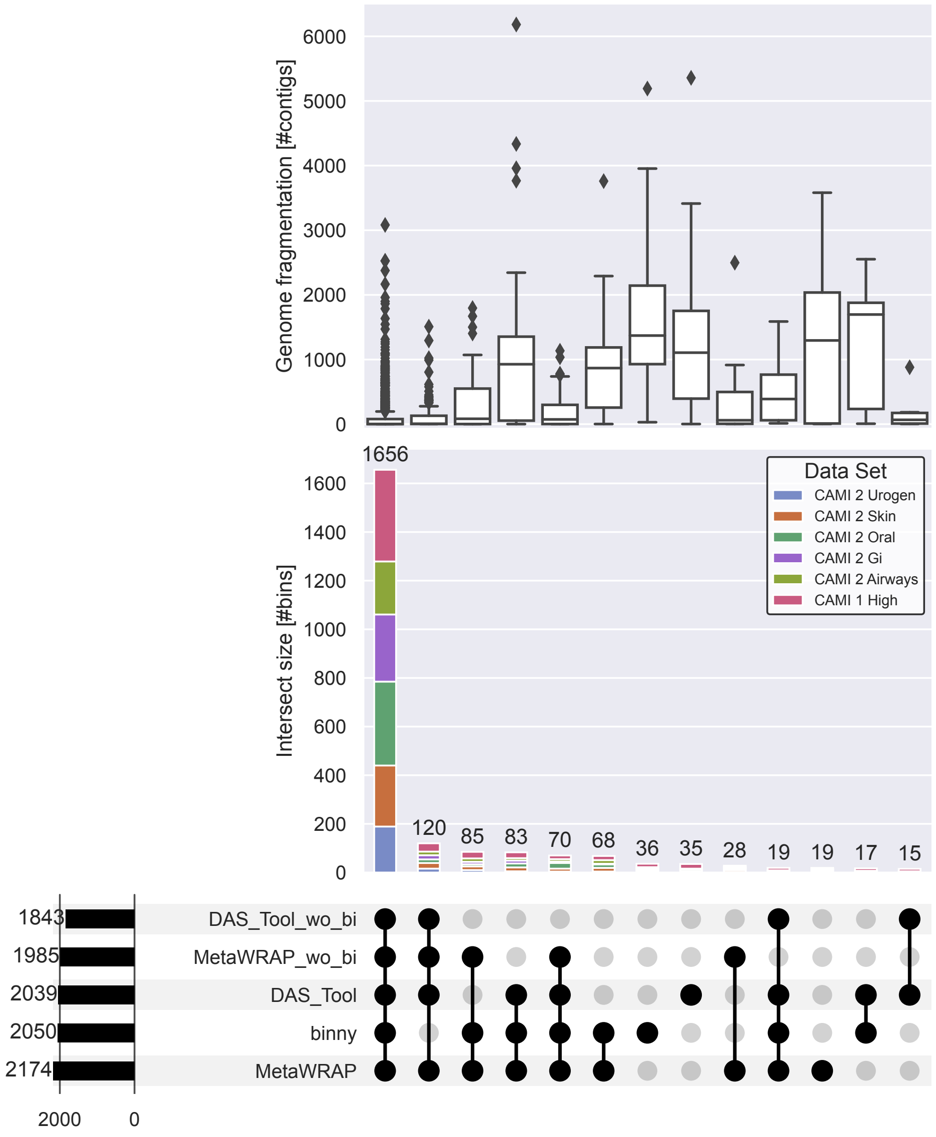


**Supplementary figure 22** UpSet plot of genome fragmentation and intersections of high quality (HQ, purity >90%, completeness >70%) MAGs of unfiltered binny and two binning refinement methods for 54 samples from six CAMI data sets. Upper panel: Genome fragmentation in number of contigs. Middle panel: Intersection size in number of HQ MAGs with proportions of MAGs stemming from the six CAMI data sets. Lower panel: Number of MAGs per binning method on the left, intersection types > 9 in the centre.

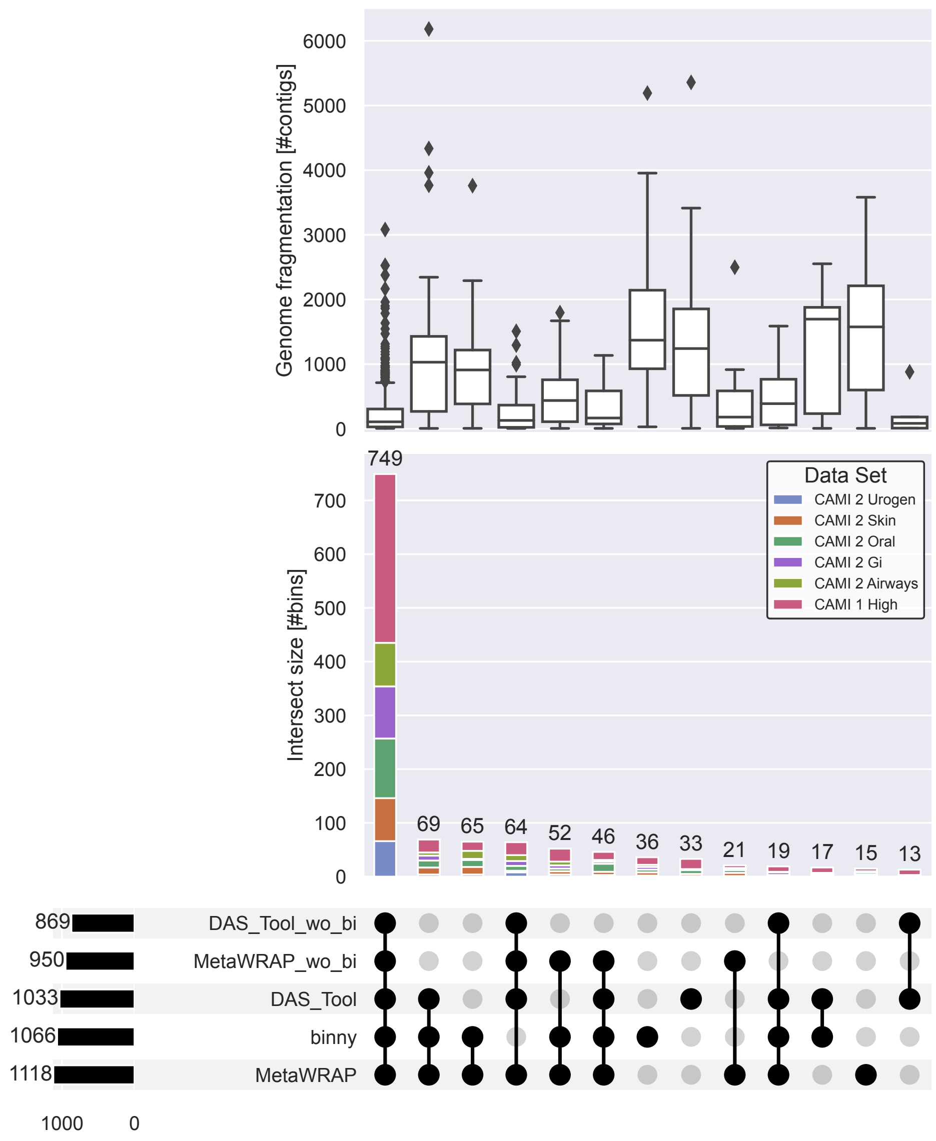


**Supplementary figure 23** UpSet plot of genome fragmentation and intersections of high quality (HQ, purity >90%, completeness >70%) MAGs of genomes consisting of more than five contigs of unfiltered binny and two binning refinement methods for 54 samples from six CAMI data sets. Upper panel: Genome fragmentation in number of contigs. Middle panel: Intersection size in number of NC MAGs with proportions of MAGs stemming from the six CAMI data sets. Lower panel: Number of MAGs per binning method on the left, intersection types > 9 in the centre.
